# Short-Term Disruption of TGFβ Signaling in Adult Mice Renders the Aorta Vulnerable to Hypertension-Induced Dissection

**DOI:** 10.1101/2024.04.22.590484

**Authors:** Bo Jiang, Pengwei Ren, Changshun He, Mo Wang, Sae-Il Murtada, Yu Chen, Abhay B. Ramachandra, Guangxin Li, Lingfeng Qin, Roland Assi, Martin A. Schwartz, Jay D. Humphrey, George Tellides

## Abstract

Hypertension and transient increases in blood pressure from extreme exertion are risk factors for aortic dissection in patients with age-related vascular degeneration or inherited connective tissue disorders. Yet, the common experimental model of angiotensin II-induced aortopathy in mice appears independent of high blood pressure as lesions do not occur in response to an alternative vasoconstrictor, norepinephrine, and are not prevented by co-treatment with a vasodilator, hydralazine. We investigated vasoconstrictor administration to adult mice 1 week after disruption of TGFβ signaling in smooth muscle cells. Norepinephrine increased blood pressure and induced aortic dissection by 7 days and even within 30 minutes that was rescued by hydralazine; results were similar with angiotensin II. Changes in regulatory contractile molecule expression were not of pathological significance. Rather, reduced synthesis of extracellular matrix yielded a vulnerable aortic phenotype by decreasing medial collagen, most dynamically type XVIII, and impairing cell-matrix adhesion. We conclude that transient and sustained increases in blood pressure cause dissection in aortas rendered vulnerable by inhibition of TGFβ-driven extracellular matrix production by smooth muscle cells. A corollary is that medial fibrosis, a frequent feature of medial degeneration, may afford some protection against aortic dissection.

## Introduction

The aorta, as the largest artery in the body subjected to the highest mechanical loads, requires the greatest mechanical strength of any blood vessel. Its complex multilaminar structure can often withstand even supra-physiological loads when constituent cells and extracellular matrix (ECM) exhibit normal organization and function. Rarely, a normal aorta will rupture when acted upon by excessive mechanical stress, as, for example, due to severe blunt trauma. More commonly, structural failure of the aortic wall occurs in response to increased blood pressure within the context of mural defects, typically age-related medial degeneration, wall thinning secondary to aortic dilatation, or inherited connective tissue disorders. A constellation of life-threatening conditions, including dissection and rupture, termed acute aortic syndrome, results when mechanical stress exceeds wall strength (1). Wall strength, however, is not measured by clinical diagnostic techniques and this lack of knowledge significantly limits our ability to predict complications (2,3). Thus, the concept of a vulnerable aorta, although challenging to quantify, deserves greater attention.

Experimental models of aortic dissection and rupture have contributed significantly to understanding aorta pathophysiology. A popular model is chronic infusion of angiotensin II (AngII) in wild-type (WT) or hyperlipidemic mice because of the simplicity of its design. Infusion of AngII at high rates increases blood pressure and, in a subset of animals, leads to aortic dilatation, dissection, or rupture (4). Yet, aortic lesions do not result after similar pressure elevations induced via infusion of an alternative vasoconstrictor, norepinephrine (NE), and AngII-mediated aortic disease is not prevented by co-treatment with a vasodilator, hydralazine (5,6). These observations have been interpreted as high blood pressure not contributing to aortic dissection and rupture, at least in this common model. An alternative explanation, however, is that pharmacologically induced hypertension does not injure aortas in the absence of underlying medial vulnerabilities and that, unlike AngII, NE does not induce sensitizing mural defects.

We, and others, are interested in the role of transforming growth factor-β (TGFβ) in aortic development, homeostasis, and disease. Pathologic variants in genes encoding TGFβ receptors, ligands, and signaling effectors associate with Loeys-Dietz syndrome, a genetic connective tissue disorder predisposing to severe aortopathy (7). Germline deletions in mice of *Tgfbr1* or *Tgfbr2*, encoding TGF-β type I and II receptors, respectively, result in embryonic lethality due to vascular defects, even when restricted to smooth muscle cell (SMC) lineages (8,9). Conditional disruption of TGFβ signaling in SMCs of the developing aorta of 3- to 6-week-old mice results in aortic thickening, dilatation, and dissection (10–12). Yet, little is known about the role of TGFβ in homeostasis of the mature aorta following completion of early postnatal growth, ECM deposition, and acquisition of SMC contractile phenotype.

In this study, we find that short-term disruption of TGFβ signaling in SMCs of adult mice predisposes to aortic dissection induced by high blood pressure within 7 days of constant infusion of NE and even within 30 minutes of boluses of NE. This vulnerable aortic phenotype results from decreased ECM production within the media and impaired cell-matrix adhesion. The rapid loss of certain collagens, such as type XVIII, within days was unanticipated in view of the far longer half-life for total collagen turnover in the normal aorta.

## Results

### Short-Term Disruption of TGFβ Signaling in SMCs of Normotensive Adult Mice does not Induce Aortopathy

We disrupted TGFβ signaling in SMCs of adult mice using a conditional gene deletion strategy that targeted both *Tgbfr1* and *Tgfbr2* to avoid possible aberrant signaling from alternate receptor pairs (13,14). Compound mutants with Cre recombinase fused to a modified estrogen receptor under control of the *Myh11* promoter enabled SMC specificity (15), while the *mT/mG* reporter allowed identification of cells with transgene recombination by green fluorescent protein (GFP) expression (16). Cre activity was induced by tamoxifen administration to 11-week-old *Tgfbr1^f/f^.Tgfbr2^f/f^.Myh11-CreER^T2^.mT/mG* mice for 5 days, hereafter designated Tgfbr1/2^iSMCKO^. Control littermates were injected with corn oil vehicle (designated as Tgfbr1/2^+/+^); other controls with intact TGFβ signaling included C57BL/6J (designated B6 WT) and tamoxifen-induced *Myh11-CreER^T2^.mT/mG* (designated GFP^iSMC^) mice. Within 7 days of starting tamoxifen (or 2 days after the last dose), 12-week-old Tgfbr1/2^iSMCKO^ mice selectively expressed GFP in most medial cells and demonstrated disrupted TGFβ signaling as evidenced by decreased phosphorylation of Smad2 (p-Smad2) in both aortas in vivo and *Myh11* lineage-marked SMCs in vitro (Figure 1A-C). Blood pressure remained unaltered based on tail-cuff measurement, ascending aorta size was unchanged on ultrasound examination, and there was no evident pathology on gross inspection or histological analysis (Figure 1D-G). In contrast, when TGFβ signaling was disrupted in 4-week-old mice via tamoxifen induction (with similar efficiency of Cre recombination in > 90% of SMCs as adult animals), 75% of the Tgfbr1/2^iSMCKO^ mice developed aneurysms, intimomedial tears, and dissection of the thoracic aorta over a 4-week period (Supplemental Figure 1), similar to previous results with deletion of *Tgfbr2* alone (10). Recalling that the aorta matures biomechanically by 8 weeks of age (17), short-term disruption of TGFβ signaling in SMCs of mature aortas does not result in overt aortic disease.

**Figure 1:**
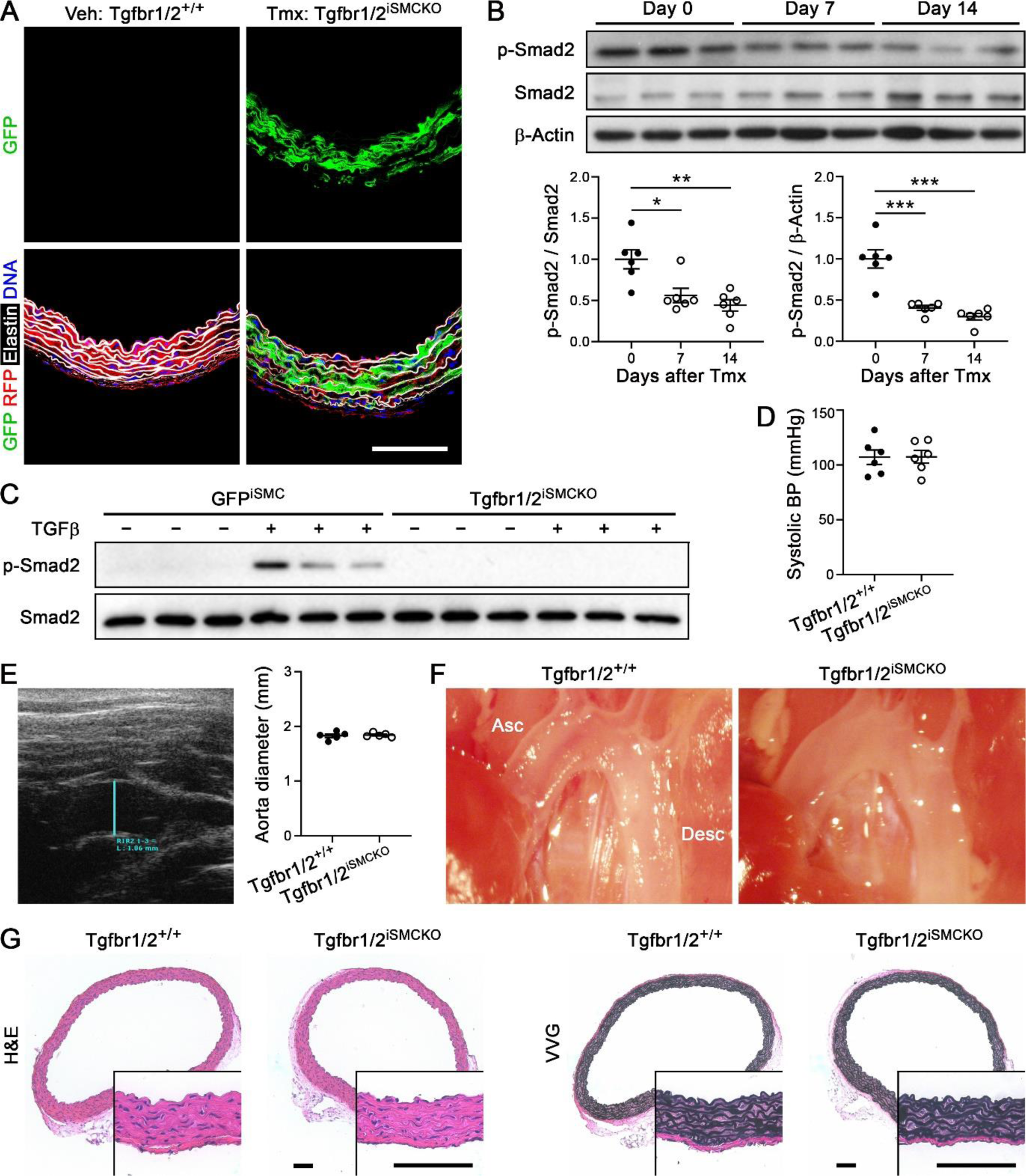
No gross phenotype 1-week after disruption of TGFβ signaling in SMCs of mature aortas. Eleven-week-old *Tgfbr1*^f/f^.*Tgfbr2*^f/f^.*Myh11*-*CreER*^T2^.*mT*/*mG* mice were injected daily with vehicle (Veh, denoted as Tgfbr1/2^+/+^) or tamoxifen (Tmx, denoted as Tgfbr1/2^iSMCKO^) for 5 days and their ascending aortas were examined at 12 weeks of age. (**A**) Expression of GFP in SMCs with Cre recombination, red fluorescent protein (RFP) in unrecombined cells, AF633 hydrazide-labeled elastin, and DAPI-labeled nuclei. (**B**) Western blots for indicated proteins at 0, 7, and 14 days after starting tamoxifen with densitometry of protein bands relative to loading controls (*n* = 6). (**C**) Similar blots of cultured SMCs isolated from tamoxifen-induced *Myh11*-*CreER*^T2^.*mT*/*mG* (denoted as GFP^iSMC^) and Tgfbr1/2^iSMCKO^ mice without or with TGFβ exposure at 1 ng/mL for 30 minutes. (**D**) Systolic blood pressure (BP) measured by tail-cuff (*n* = 6). (**E**) Ultrasound examination of ascending aorta diameter (blue line, *n* = 5). (**F**) In situ examination of ascending (Asc) and descending (Desc) thoracic aortas with unremarkable appearances. (**G**) Hematoxylin and eosin (H&E) and Verhoeff–Van Gieson (VVG) stains. Scale bars: 100 μm. Data are shown as individual values with mean ± SEM; \**P* < 0.05, \*\**P* < 0.01, \*\*\**P* < 0.001 by t-test (D, E) or 1-way ANOVA with Tukey’s multiple comparisons test (B).

### Continuous Infusion of NE Induces Aortic Dissection in Adult Tgfbr1/2^iSMCKO^ Mice

We investigated if NE infusion exposes underlying structural vulnerabilities by infusing 12-week-old Tgfbr1/2^iSMCKO^ mice at a pressor dose known not to induce injury of non-vulnerable aortas (5,6). A relatively short infusion duration of 7 days was implemented because mild hemorrhagic lesions of the aorta can resolve with few visible sequelae after the longer durations commonly employed in experimental studies (e.g., 28 days) and dissections rarely occur late during this extended period (6,18). Continuous infusion of NE at 3.88 μg/kg/min for 7 days using a subcutaneous (s.c.) osmotic minipump increased blood pressure, but to a lesser degree than AngII infusion at the standard dose of 1 μg/kg/min (Figure 2A,B). Examination under a dissecting microscope after saline flush of luminal blood revealed focal hematomas of the ascending aorta and aortic arch, occasionally extending into the descending thoracic aorta, in 5.6% of saline-, 51.1% of NE-, and 100% of AngII-infused Tgfbr1/2^iSMCKO^ mice (Figure 2C,D). All 81 saline- and NE-infused animals survived, whereas 3 of 21 AngII-infused animals died between 3-6 days and were confirmed to have ruptured descending thoracic aortas with accumulation of intracavitary blood (Supplemental Figure 2A,B). Histological analysis verified the hemorrhagic lesions of ascending aortas as dissections, defined as blood extravasation into the media (Figure 2E). Yet not all intimomedial tears visible by histology associated locally with mural hematomas or medial dissection (Supplemental Figure 2C). We confirmed that NE infusion for 7 days did not result in hemorrhagic lesions in 12-week-old B6 WT mice (Supplemental Figure 2D). In summary, NE-induced hypertension caused aortopathy in adult Tgfbr1/2^iSMCKO^ mice but not in age- and sex-matched mice with intact TGFβ signaling.

**Figure 2:**
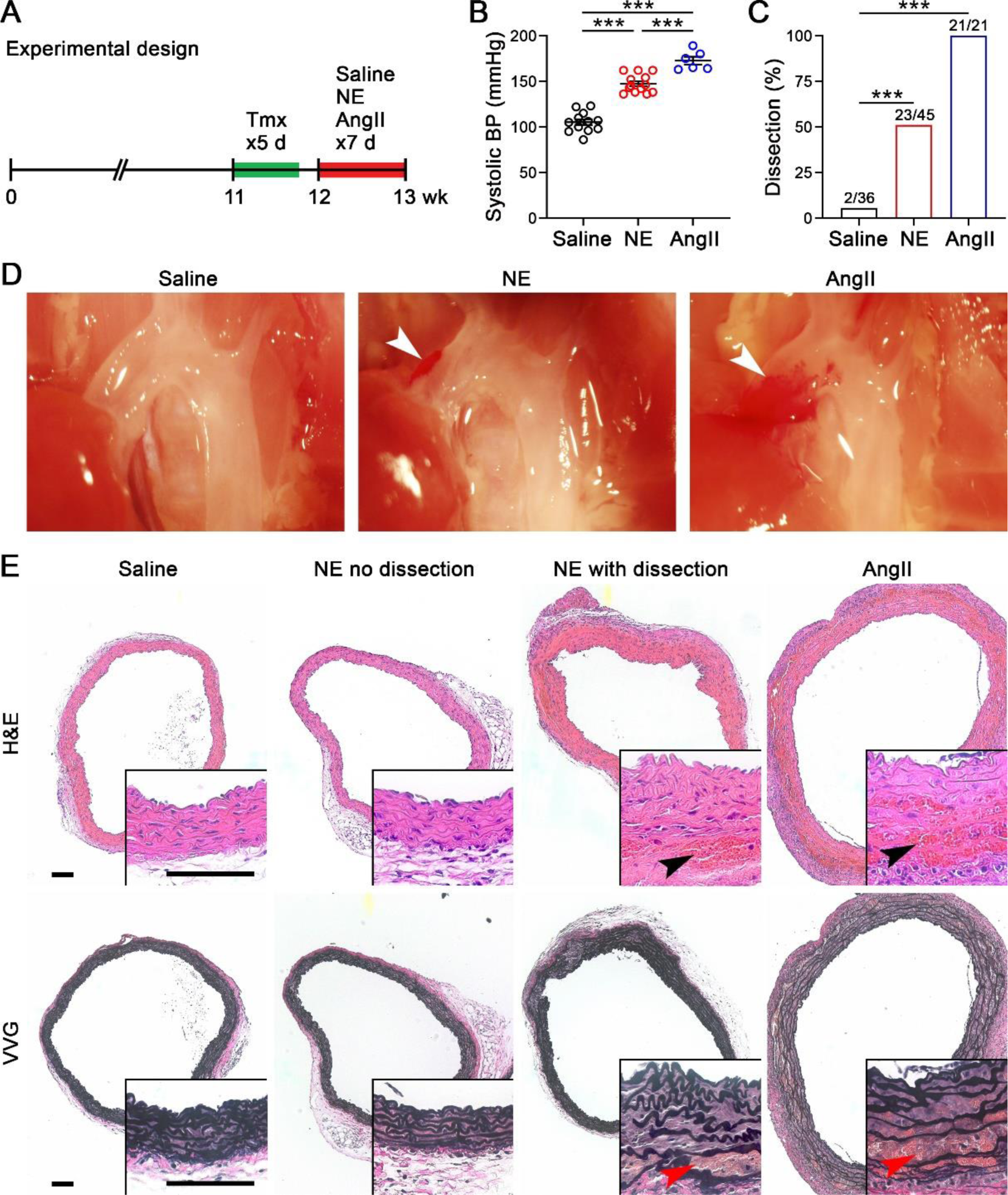
Continuous 1-week infusion of NE induces dissection in vulnerable aortas. (**A**) One week after starting tamoxifen (Tmx), namely 2 days after the last dose, 12-week-old Tgfbr1/2^iSMCKO^ mice were infused with saline, NE at 3.88 μg/kg/min, or AngII at 1 μg/kg/min by osmotic minipump for 7 days and examined at 13 weeks of age. (**B**) Systolic blood pressure (BP) measured by Millar catheter in control and vasoconstrictor-infused animals (*n* = 6-12). (**C**) Incidence of aortic dissection in saline- (*n* = 2/36), NE- (*n* = 23/45), and AngII- (*n* = 21/21) infused animals. (**D**) In situ examination by dissecting microscope after saline flush via the left ventricle showing mural hematomas of the ascending aorta (arrows). (**E**) Hematoxylin and eosin (H&E) and Verhoeff–Van Gieson (VVG) stains confirmed aortic dissections by blood extravasation into the media (arrows) in a subset of NE- and all AngII-infused animals. Scale bars: 100 μm. Data are shown as individual values with mean ± SEM, \*\*\**P* < 0.001 by 1-way ANOVA with Tukey’s multiple comparisons test (B) or Fisher’s exact test between study groups versus control (C).

### Immediate Aortic Dissection after NE Bolus in Adult Tgfbr1/2^iSMCKO^ Mice

To determine if short-term disruption of TGFβ signaling was sufficient for aortic dissection triggered by elevated blood pressure or if NE-mediated remodeling of the aorta was also necessary for disease susceptibility, the duration of NE infusion was reduced in steps. In pilot experiments, hematomas occurred in 2 of 3 aortas at 2 days, 3 of 4 aortas at 1 day, and 3 of 4 aortas at 12 hours of NE infusion. We then tested the effects of single intraperitoneal (i.p.) injections of NE at a dose of 1.28 mg/kg known to elevate blood pressure acutely in mice (19). Central blood pressure monitoring by invasive catheter via the right carotid artery revealed a rapid increase of blood pressure, peaking within 5 minutes and remaining elevated over 30 minutes though at lower levels than that induced by single intraperitoneal injection of AngII at a pressor dose of 0.64 mg/kg (19) (Figure 3A,B, Supplemental Figure 3A,B). Examination under a dissecting microscope revealed hematomas of the ascending aorta and/or aortic arch in 0% of saline-, 52.9% of NE-, and 69.2% of AngII-infused Tgfbr1/2^iSMCKO^ mice after 30 minutes (Figure 3C,D). The hemorrhagic lesions were limited in extent and there were no immediate deaths after boluses of vasoconstrictors. Histology confirmed medial dissection (Figure 3E). Concomitant treatment with a vasodilator, hydralazine at 10 mg/kg i.p., prevented NE-induced pressure elevations and aortic dissections over 30 minutes (Supplemental Figure 3C-E). Hydralazine, however, did not block blood pressure responses to AngII (possibly because of greater vasoconstrictor responses at the doses tested) and effects on aortic dissection by this combination of agents were not investigated (Supplemental Figure 3F). Thus, short-term disruption of TGFβ signaling was sufficient to render the aortas vulnerable to dissection induced by high blood pressure independent of possible NE-mediated aortic remodeling.

**Figure 3:**
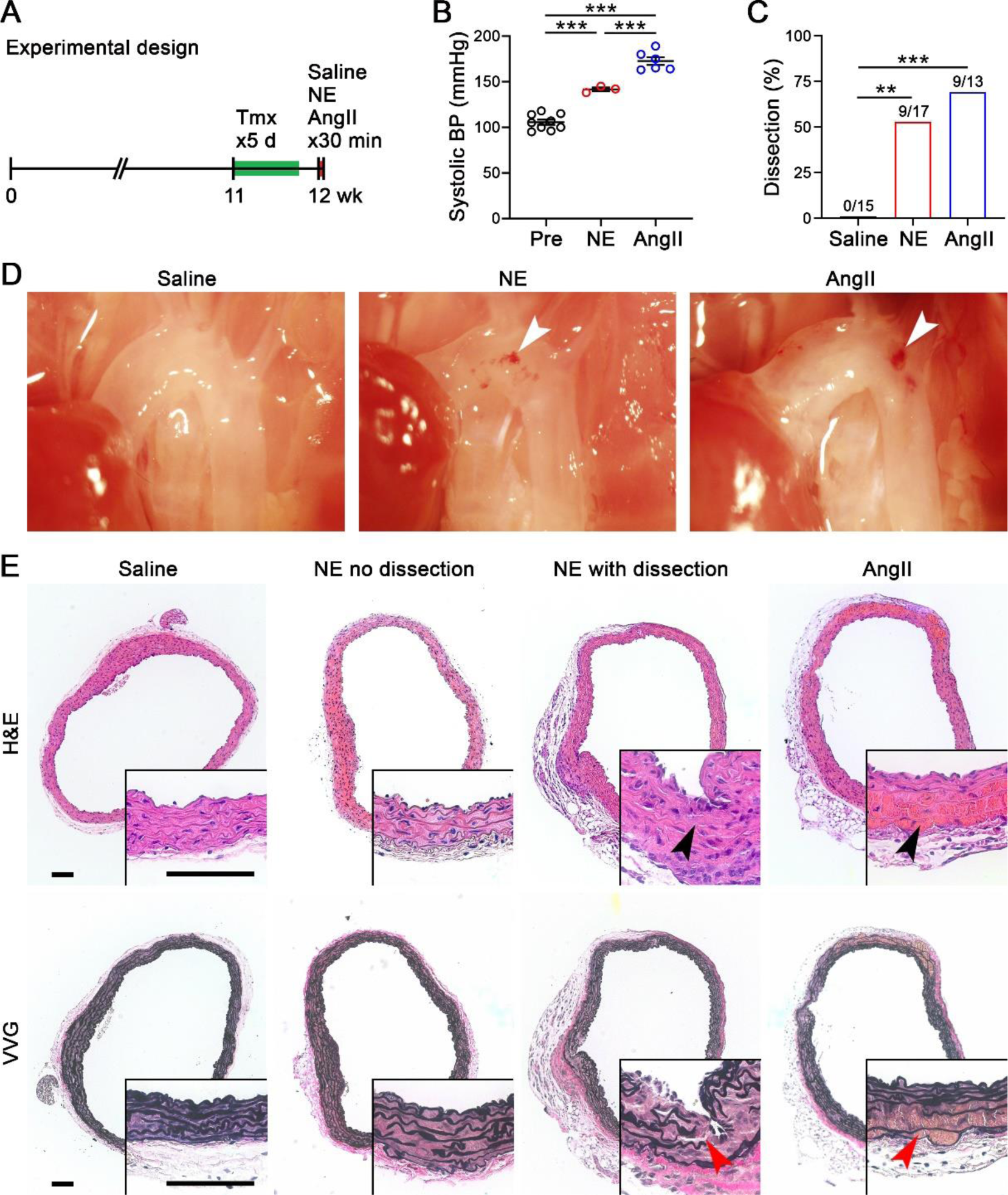
Bolus of NE induces dissection in vulnerable aortas. (**A**) One week after starting tamoxifen (Tmx), or 2 days after the last dose, 12-week-old Tgfbr1/2^iSMCKO^ mice were injected intraperitoneally with saline, NE at 1.28 mg/kg, or AngII at 0.64 mg/kg and examined after 30 minutes. (**B**) Maximum systolic blood pressure (BP) measured by Millar catheter before (Pre) and 30 minutes after injection (*n* = 3-6). (**C**) Incidence of aortic dissection in saline- (*n* = 0/15), NE- (*n* = 9/17), and AngII- (*n* = 9/13) injected animals. (**D**) In situ examination by dissecting microscope after saline flush via the left ventricle showing mural hematomas of the ascending aorta (arrows). (**E**) Hematoxylin and eosin (H&E) and Verhoeff–Van Gieson (VVG) stains showing medial dissections (arrows) in a subset of NE- and AngII-injected mice. Scale bars: 100 μm. Data are shown as individual values with mean ± SEM bars. \*\**P* < 0.01, \*\*\**P* < 0.001 by 1-way ANOVA with Tukey’s multiple comparisons test (B) or Fisher’s exact test between vasoconstrictors versus control (C).

### Altered Traction on and Fragmentation of SMCs with Increasing Medial Delamination

Dissected aortas after vasoconstrictor infusion for 30 minutes were examined further to define immediate histopathological outcomes prior to time-dependent responses to injury, such as apoptosis, inflammation, and fibrosis. Specimens were perfusion-fixed at physiological pressure to straighten the medial laminae (Supplemental Figure 4); because this process could cause distension artifacts, we examined both pressure-fixed and unpressurized specimens. Partial tears through the intima and subjacent media marked by thrombus served as entry sites for blood extravasation into the media, but an intact external elastic lamina and adventitia prevented free rupture (Supplemental Figure 5A,B). Confocal studies with reagents specific for SMC cytoskeleton, red blood cells (RBCs), and elastin revealed disease heterogeneity with blood tracking circumferentially and axially from intimomedial tears along the outer laminae while sparing inner laminae (Figure 4A-E). Non-widened laminae consisted of tightly packed SMCs bordered by elastic fibers. In moderately widened laminae there was separation of neighboring SMCs from each other with intervening RBCs; the long axis of SMCs changed from circumferential to radial with persistent attachment to ill-defined extensions of elastic laminae (intralaminar elastic fibers). In contrast, markedly widened laminae were filled with RBCs amid infrequent SMC fragments. Similar heterogenous effects were found with alternative reagents labeling SMC plasma membrane and collagen: these components lined elastic fibers in areas without dissection, individual or small groups of RBCs infiltrated between SMCs and fibrillar matrix in laminae with mild dissection, and SMC membrane fragments were adherent to elastic fibers in regions with marked delamination (Figure 4F-J). Transmission electron microscopy confirmed displacement of SMCs by RBCs with residual cell and matrix fragments attached to widened laminae, but an unremarkable relationship of SMCs to elastic and collagen fibers in adjacent non-widened laminae (Figure 4K,L). In contrast to hemorrhagic aortic lesions, circumferentially-oriented SMCs were uniformly attached to ECM and neighboring cells in aortas prior to NE administration or without dissection despite pharmacologically induced pressure elevation (Supplemental Figure 5C,D). The association of varying medial delamination with RBC extravasation and SMC fragmentation suggests underlying TGFβ-dependent defects in cell-cell or cell-matrix adhesion as well as contractility or cytoskeletal strength, respectively.

**Figure 4:**
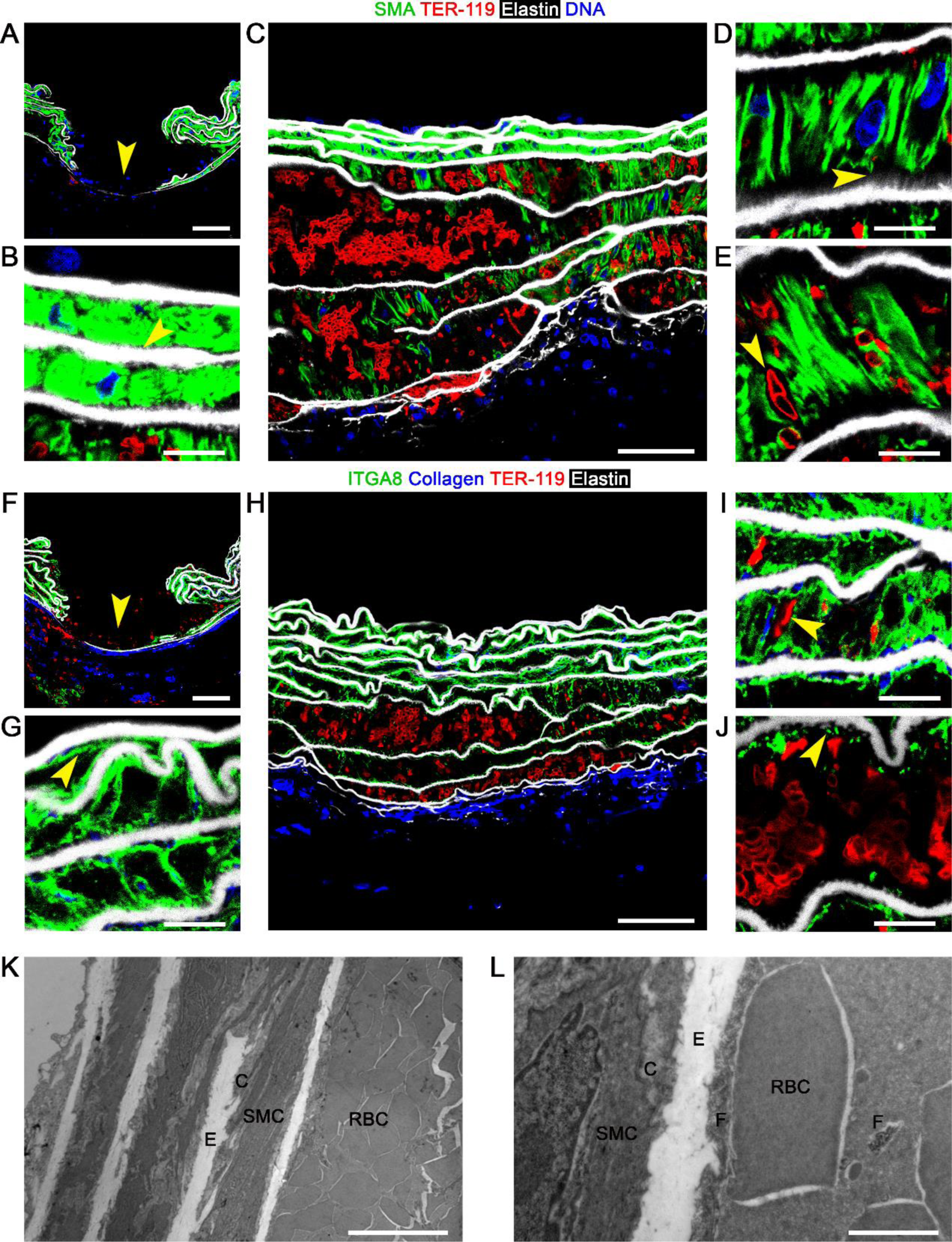
Traction on and rapid fragmentation of SMCs. Twelve-week-old Tgfbr1/2^iSMCKO^ mice were infused with NE at 1.28 mg/kg i.p. for 30 minutes and the ascending aortas examined. Confocal microscopy after labeling smooth muscle α-actin (SMA) for SMC cytoskeleton (green), TER-119 for RBCs (red), AF633 hydrazide for elastin (white), and DAPI for nuclei (blue) shows (**A**) intimomedial entry tear (arrow), (**B**) non-widened inner laminae with SMCs adjacent to elastic fibers (arrow), (**C**) varying RBC accumulation in outer laminae, (**D**) widened laminae with radially-oriented SMCs attached to ill-defined intralaminar elastic fibers (arrow), and (**E**) RBCs between SMCs (arrow). Alternative labeling to integrin α8 (ITGA8) for SMC plasma membrane (green) and CNA35 to collagen (blue) shows (**F**) intimomedial entry tear (arrow), (**G**) non-widened laminae with intact SMC plasma membranes (arrow), (**H**) varying RBC accumulation in outer laminae, (**I**) RBCs among SMCs (arrow), and (**J**) widened lamina with attached SMC plasma membrane fragments (arrow) and areas where elastic laminae are stripped clean of cell and fibrillar matrix. Transmission electron microscopy showing (**K**) RBC accumulation in outer laminae and (**L**) non-widened lamina with SMCs contacting elastic (E) and collagen (C) fibers adjacent to widened lamina with RBCs abutting elastic and collagen fibers and cellular fragments (F). Pressure-fixed (A, F, K, L) and unpressurized (B-E, G-J) specimens. Scale bars: 50 μm (A, C, F, H), 10 μm (B, D, E, G, I-K), and 2 μm (L).

### Limited Biomechanical Impairment 1 Week After Disrupting TGFβ Signaling in Mature Aortas

We investigated if vulnerability of the aortic wall following TGFβ disruption and 7 days of NE infusion was related to changes in bulk structural or material properties. Passive testing (i.e., without SMC contractility) of ascending aortas was performed ex vivo using a computer-controlled multiaxial device (20). The vessel segments did not leak when extended axially and pressurized over physiological ranges, implying overall structural integrity. Most biomechanical metrics remained similar despite the presence of small dissections in a subset of aortas from NE-infused animals (Figure 5A,B). A modest decrease in axial stretch and distensibility of aortas from NE-infused Tgfbr1/2^iSMCKO^ mice suggested modest elastic fiber breaks or accumulation of non-elastic material such as glycosaminoglycans, blood, or debris. Elastic energy storage, a key function of the aortic wall stemming primarily from intact elastic laminae, was yet unchanged. The minimal differences in mechanical properties were also illustrated by overlapping pressure-radius (structural behavior) and stress-stretch (material behavior) curves (Figure 5C). Active testing of SMC contractility revealed decreased ex vivo responses to phenylephrine in ascending aortas of NE-infused Tgfbr1/2^iSMCKO^ mice suggesting desensitization to adrenergic stimuli rather than global impairment in contractile function as KCl responses were not affected (Figure 5D,E). Although biomechanical properties averaged over an entire vessel segment showed only small differences after short-term disruption of TGFβ signaling, this does not exclude local micro-defects with a potential to initiate and propagate aortic dissection in response to a hypertensive stimulus.

**Figure 5.**
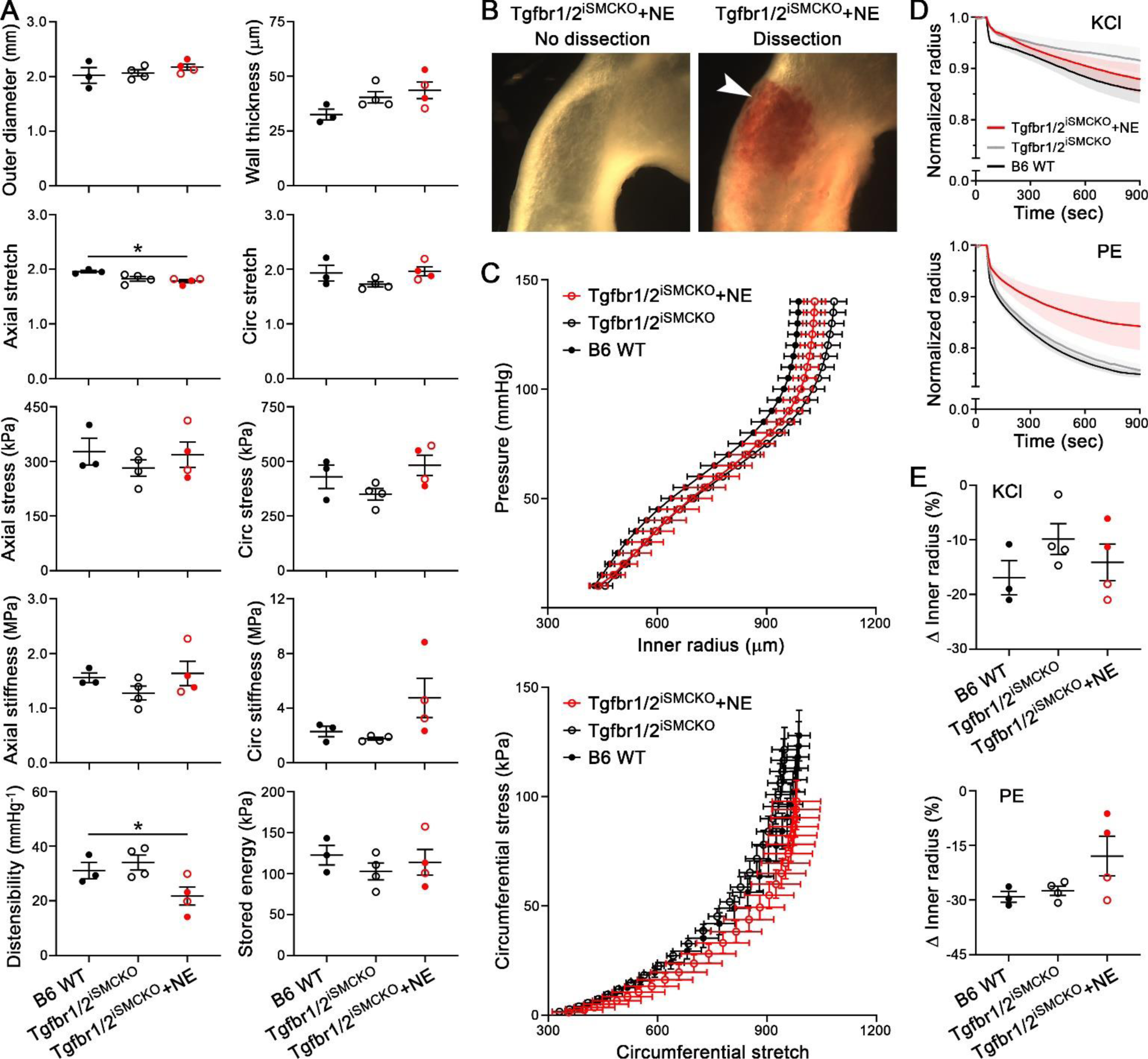
Limited impairment of bulk biomechanical properties 1 week after disrupting TGFβ signaling in mature aortas. Vessel-level biomechanical testing was performed on a sub-set of ascending aortas from untreated B6 WT mice and untreated or 7-day NE-infused Tgfbr1/2^iSMCKO^ mice. (**A**) Outer diameter, wall thickness, axial and circumferential (circ) stretch, mean wall stress, and material stiffness, distensibility, and stored energy at group-specific systolic pressures. (**B**) Unpressurized ascending aortas from NE-infused Tgfbr1/2^iSMCKO^ mice without or with dissection (arrow). (**C**) Overlapping pressure-radius and circumferential stress-stretch curves among the 3 groups. (**D**) Vasoconstriction, against a fixed pressure at the in vivo axial stretch, responses to KCl and phenylephrine (PE) assessed by reduction of normalized inner radius over time. (**E**) Steady state change of inner radius in response to KCl and PE. Data are shown as individual values with mean ± SEM bars (A, E) or means ± SEM with connecting lines (B, D). *n* = 3-4 per group, NE infusion resulted in no dissection (open red symbols, *n* = 2) or dissection (filled red symbols, *n* = 2). \**P* < 0.05, 1-way ANOVA with Tukey’s multiple comparisons test for steady state values (A, E).

### Disrupted TGFβ Signaling Alters Transcription of ECM and Regulatory Contractile Molecules

To identify transcriptional responses associated with short-term disruption of TGFβ signaling and vulnerability to hemodynamic stresses, we performed whole transcriptome profiling. Bulk RNA sequencing (RNA-seq) of thoracic aortas from 12-week-old Tgfbr1/2^iSMCKO^ versus B6 WT mice revealed 840 differentially expressed genes with > 2-fold change, including an ECM (*Col15a1*) and a regulatory contractile (*Mylk4*) molecule as the most significantly downregulated transcripts (Supplemental Figure 6A). Gene ontology profiling identified cell adhesion and ECM among over-represented terms (Supplemental Figure 6B). Since analysis of whole aortic tissue may be confounded by transcript expression from non-recombined SMCs and cell types other than SMCs, we analyzed isolated cells after cryophilic enzyme digestion at 4 °C to prevent cell stress artifact (21). Bulk RNA-seq of *Myh11* lineage-marked SMCs from 12-week-old Tgfbr1/2^iSMCKO^ versus GFP^iSMC^ mice showed 597 differentially expressed genes, of which most were downregulated, including those for diverse ECM components – fibrillar (e.g., *Col1a2*, *Col3a1*) and non-fibrillar (e.g., *Col15a1*, *Col18a1*) collagens, elastic fibers and microfibrils (e.g., *Eln*, *Mfap4*), basement membrane (e.g., *Col4a2*, *Col4a5*), enzymes for ECM organization (e.g., *Lox*), matricellular molecules (e.g., *Ccn2*), and various proteoglycans (e.g., *Bgn*) – as well as regulatory contractile molecules (e.g., *Mylk4*) (Figure 6A). Gene ontology enrichment analysis also identified ECM and cell adhesion themes predominating among over-represented terms (Figure 6B). Changes in gene expression were similar in SMCs from ascending/arch and descending thoracic segments (Supplemental Figure 6C). The differential expression of genes in isolated SMCs only partially overlapped with that of aortas, however, because the expression of many ECM molecules (but not *Col15a1* and *Col18a1*) predominated in fibroblasts and thus confounded whole tissue analysis (Supplemental Figure 6D-F). Single-cell RNA-seq analysis of *Myh11* lineage-marked SMCs from ascending/arch segments revealed a primary partition of populations by TGFβ signaling state and further clustering of cells within each experimental condition (Figure 6C,D). Gene expression of selected ECM and regulatory contractile molecules, quantified as differentially expressed by bulk RNA-seq, also showed clear separation by experimental condition in single-cell RNA-seq analysis demonstrating ubiquitous effects of TGFβ signaling disruption among SMC subpopulations (Figure 6E). The predominant loss of transcripts in SMCs from Tgfbr1/2^iSMCKO^ versus GFP^iSMC^ mice was unlikely due to dying cells as cell viability and RNA quality control metrics were similar between experimental groups and replicates (Supplemental Figure 7). Together, these unbiased analyses document rapid downregulation of genes for many ECM molecules secreted by SMCs, likely affecting cell-matrix interactions with relatively few changes in contractile molecules.

**Figure 6:**
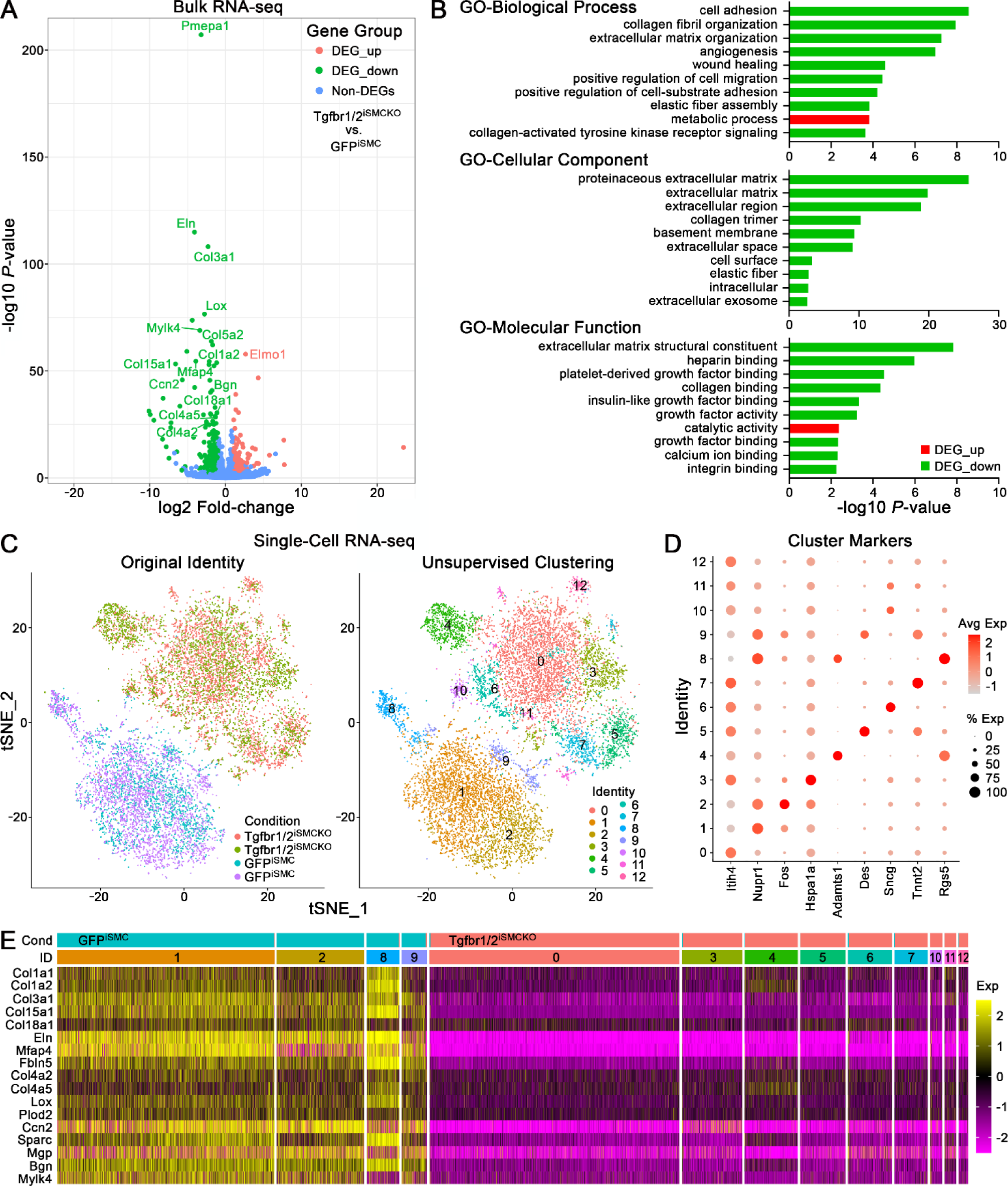
Whole transcriptome profiling 1 week after disrupting TGFβ signaling in SMCs of mature aortas. *Myh11* lineage marked SMCs were isolated from thoracic aortas of 12-week-old GFP^iSMC^ and Tgfbr1/2^iSMCKO^ mice. Bulk RNA-seq (*n* = 12) shown as (**A**) volcano plot of differentially expressed genes (DEG) and (**B**) gene ontology (GO) enrichment analysis. Single cell RNA-seq (*n* = 4) shown as (**C**) t-distributed Stochastic Neighbor Embedding (tSNE) plots with 13 clusters identified and cells distinctly partitioned by experimental condition, (**D**) dot plot of log2 average expression and % expression of cluster markers (markers for 4 clusters containing <3% of total cells are not shown), and (**E**) heatmap of selected ECM and regulatory contractile molecules (identified as differentially expressed in bulk RNA-seq analysis) illustrating uniformly decreased expression in almost all cells with TGFβ signaling disruption irrespective of SMC subclusters (clusters 0, 3-7, 10-12).

### Altered Expression of Regulatory Contractile Molecules do not Contribute to Aortic Dissection after Short-Term TGFβ Signaling Disruption in Adult Tgfbr1/2^iSMCKO^ Mice

We initially considered if changes in SMC contractile or synthetic phenotypes contributed to aortic vulnerability as transcripts involved in these functions were among the top differentially expressed genes by RNA-seq of aortas and SMCs. We examined the role of *Mylk4* encoding myosin light chain kinase-4 because maximal SMC contraction prevents medial delamination ex vivo (22). Quantitative RT-PCR analysis of whole aortic tissue confirmed decreased *Mylk4*, although *Mylk* encoding smooth muscle myosin light chain kinase, the dominant isoform of the enzyme in SMCs, was increased corresponding to bulk RNA-seq analysis (Figure 7A,B). As expected, *Mylk2* and *Mylk3*, encoding isoforms expressed in skeletal and cardiac muscle, respectively, were not detected in SMCs. Western blot analysis showed persistent activation of myosin light chain-2 (the target of myosin light chain kinases), suggesting that downregulation of *Mylk4* was not of functional significance (Figure 7C-E). We further tested if inhibition of SMC contraction potentiated effects of TGFβ signaling disruption using various pharmacological agents as a gain-of-function (more aptly gain-of-dysfunction) approach (Figure 7F). Inhibition of myosin light chain kinase by ML-7 or indirectly by a calcium channel blocker, nifedipine, or a Rho kinase inhibitor, Fasudil, did not significantly alter the incidence of NE-triggered aortic dissection, though with a confounding lowering of blood pressure by the former 2 agents (Figure 7G,H). The limited data with at least 1 myosin light chain modulator in the context of persistently increased blood pressure from NE suggest that changes in SMC contractility are not likely to cause the dissection phenotype in this model of TGFβ signaling disruption.

**Figure 7:**
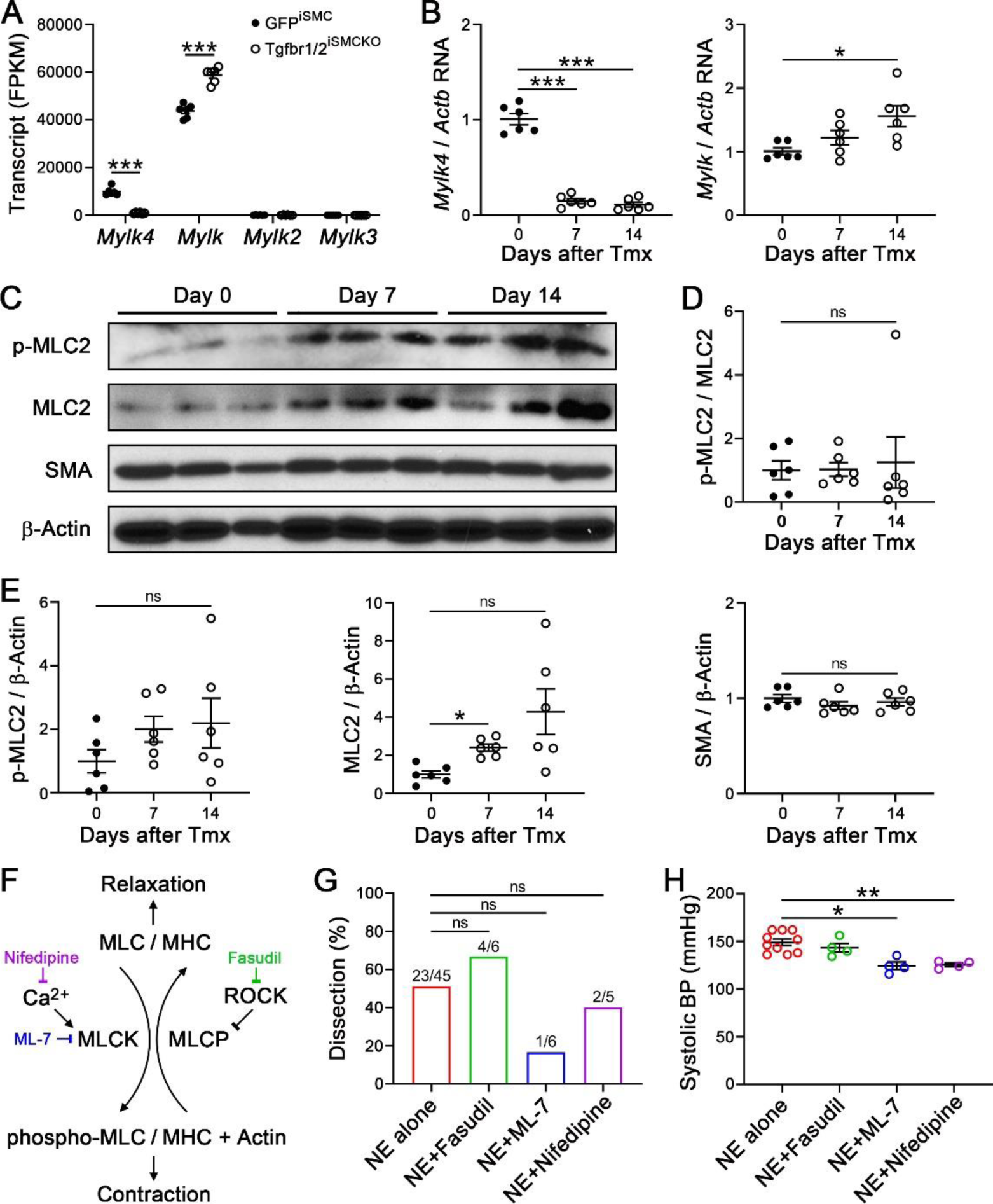
Altered expression of regulatory contractile molecules do not contribute to aortic dissection after disrupting TGFβ signaling. Thoracic aortas were analyzed after various imposed conditions. (**A**) Bulk RNA-seq for *Mylk4*, *Mylk*, *Mylk2*, and *Mylk3* as fragments per kilobase million (FPKM) from GFP^iSMC^ and Tgfbr1/2^iSMCKO^ mice (*n* = 6). (**B**) Quantitative RT-PCR for *Mylk4* and *Mylk*, relative to *Actb*, in aortas of Tgfbr1/2^iSMCKO^ mice at 0-14 days after starting tamoxifen (Tmx, *n* = 6). (**C**) Western blots for phospho-myosin light chain-2 (p-MLC2), myosin light chain-2 (MLC2), smooth muscle α-actin (SMA) as a SMC marker, and β-actin as loading control in aortas of Tgfbr1/2^iSMCKO^ mice at 0-14 days after tamoxifen. Densitometry of protein bands relative to (**D**) MLC2 or (**E**) β-actin (*n* = 6). (**F**) Phosphorylation of regulatory myosin light chain (MLC) by myosin light chain kinase (MLCK) leads to myosin heavy chain (MHC)-actin mediated contraction, whereas dephosphorylation by myosin light chain phosphatase (MLCP) enables relaxation. Vasoconstrictors activate MLCK via Ca^2+^ or inhibit MLCP via Rho-Rho kinase (ROCK); phospho-MLC and SMC contractility are inhibited by the MLCK inhibitor, ML-7, the Ca^2+^ channel blocker, Nifedipine, and the ROCK inhibitor, Fasudil. (**G**) Incidence of aortic dissection in Tgfbr1/2^iSMCKO^ mice infused with NE alone (*n* = 23/45) or with concomitant treatment with Fasudil (*n* = 4/6), ML-7 (*n* = 1/6), or Nifedipine (*n* = 2/5) for 7 days. (**H**) Systolic blood pressure (BP) measured by tail-cuff after NE-infusion ± pharmacological agents for 7 days (*n* = 4-10). Data are shown as individual values with mean ± SEM bars. \**P* < 0.05, \*\**P* < 0.01, \*\*\**P* < 0.001, or not significant (ns) by 2-way ANOVA with Sidak’s multiple comparisons test (A), 1-way ANOVA with Tukey’s multiple comparisons test (B, D, E, H), or Fisher’s exact test between combined treatments vs. NE alone (G).

### Decreased Medial Collagen Following Short-Term TGFβ Signaling Disruption in SMCs of Mature Aortas

Based on the whole transcriptome findings, we further investigated the expression of ECM molecules synthesized by SMCs. Quantitative RT-PCR revealed a rapid decline in *Col15a1* and *Col18a1* in thoracic aortas within 7 days of TGFβ receptor deletion (relevant for 30-minute vasoconstrictor experiments) and similarly at 14 days (relevant for 7-day vasoconstrictor experiments) (Figure 8A). Other collagen transcripts were not measured because of the confounding influence of the adventitia. Although the greater collagen content of the adventitia overshadows that in other vascular wall compartments, medial collagen protein is detected after overexposure of microscopy images (23). Medial collagen appeared decreased by confocal microscopy using a pan collagen-binding probe and was confirmed as modestly reduced by quantitative analysis of histological stains (Figure 8B-D). Assessment of individual collagens for which suitable reagents were available showed varying results, including substantially less collagen XVIII but less reduction in collagen I and III (Figure 8E, Supplemental Figure 8A,B). In keeping with SMC specificity of *Col18a1* by single-cell RNA-seq analysis, collagen XVIII was not detected in the adventitia except in covering mesothelial cells (visceral pericardium). Collagen XVIII co-localized with collagen IV, but not collagen I, suggesting interactions with SMC basement membrane (Figure 8F). Similar co-localization was seen with another basement membrane component, perlecan (Supplemental Figure 8C,D). Collagen IV and perlecan encircled SMCs but did not appear decreased within 7 days of TGFβ receptor deletion (Supplemental Figure 8E,F). Decreased synthesis of ECM molecules was associated with phenotypic modulation of SMCs to a less synthetic state as evidenced by a decreased proportion of endoplasmic reticulum relative to contractile filament markers (Figure 8G). Thus, certain medial collagens decreased within 7 days of disrupting TGFβ signaling in SMCs under physiological hemodynamic conditions.

**Figure 8:**
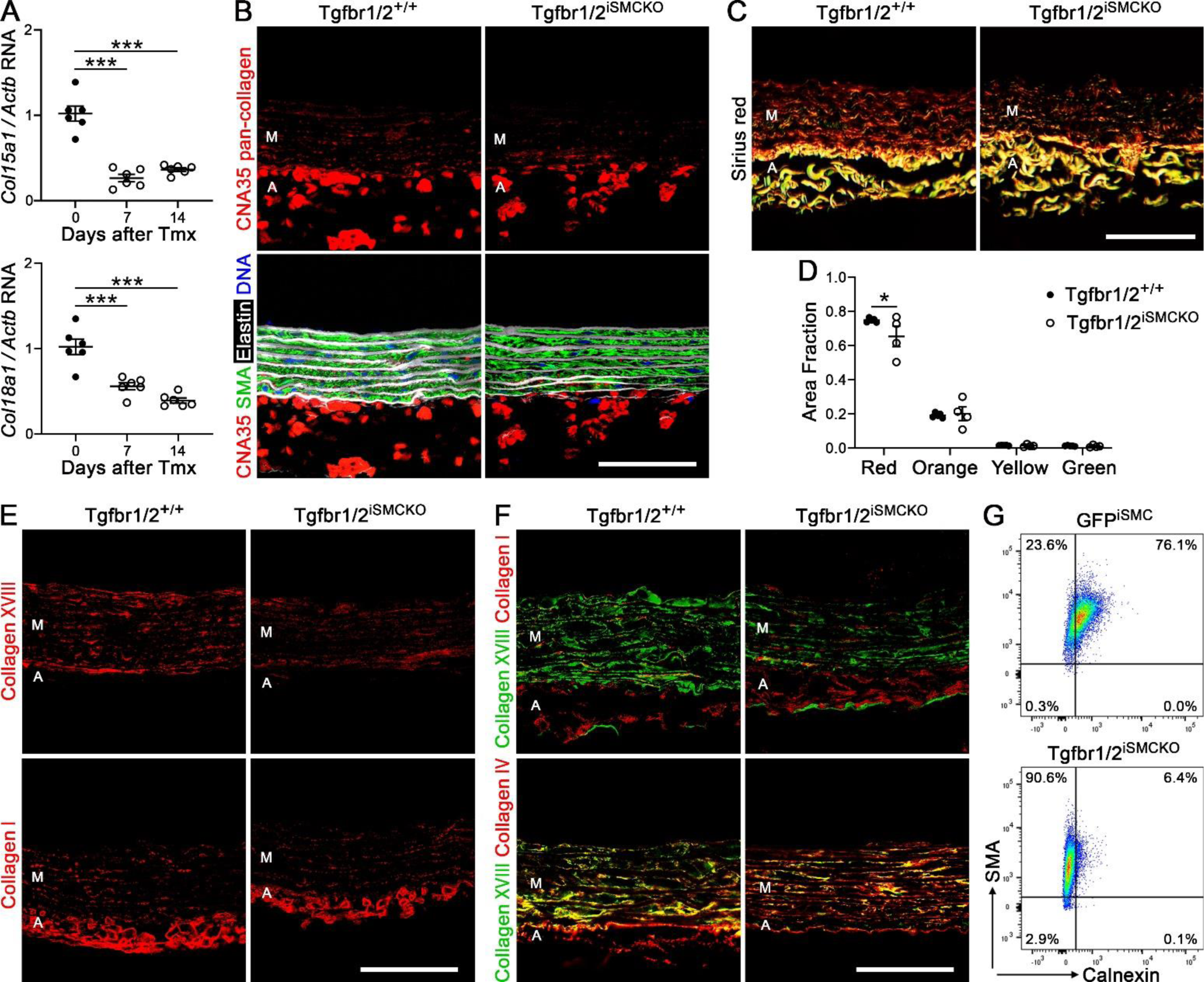
Decreased medial collagen 1 week after disrupting TGFβ signaling in SMCs of mature aortas. Thoracic aortas were analyzed after various imposed conditions. (**A**) Quantitative RT-PCR for *Col15a1* and *Col18a1* transcripts, relative to *Actb*, at 0-14 days after starting tamoxifen (Tmx, *n* = 6). Collagen protein studies were performed in 12-week-old Tgfbr1/2^+/+^ and Tgfbr1/2^iSMCKO^ mice at 7 days after starting tamoxifen. (**B**) Confocal microscopy after labeling with the collagen-binding probe CNA35 (red) alone or overlaid with smooth muscle α-actin (SMA) for SMCs (green), AF633 hydrazide for elastin (white), and DAPI for nuclei (blue). (**C**) Picrosirius red staining for collagen under polarized light. (**D**) Quantification of picrosirius red polarized colors as fraction of media area (*n* = 4-5). (**E**) Presence of collagen XVIII (red) or collagen I (red). (**F**) Overlay of collagen XVIII (green) and I (red) or collagen XVIII (green) and IV (red) demonstrating co-localization (yellow). (**G**) Intracellular presence of the contractile filament marker SMA versus the endoplasmic reticulum marker calnexin by flow cytometry consistent with reduction in synthetic state of SMCs from Tgfbr1/2^iSMCKO^ vs. GFP^iSMC^ mice. Formalin-fixed, paraffin-embedded sections (B, C, E) and frozen, OCT-embedded sections (F) of pressure-fixed (B, E, F) and unpressurized (C) specimens. The media (M) and adventitia (A) are identified by the presence or absence of elastic laminae. Scale bars: 50 μm. Data are shown as individual values with mean ± SEM bars. \**P* < 0.05 and \*\*\**P* < 0.001 by 1-way ANOVA with Tukey’s multiple comparisons test (A) and 2-way ANOVA with Sidak’s multiple comparisons test (D).

### Impaired ECM Predisposes to Aortic Dissection in Adult Tgfbr1/2^iSMCKO^ Mice

To determine if decreased deposition of functional collagen contributes to a vulnerable aorta phenotype, we administered β-aminopropionitrile (BAPN), an inhibitor of newly synthesized collagen cross-linking, at 150 mg/kg/d s.c. for 14 days via continuous infusion. This dose of BAPN is known not to induce overt aortopathy in adult B6 WT mice when administered from 9 to 11 weeks of age (24) but can impair ECM integrity due to constituents that turnover rapidly, thus enabling another gain-of-dysfunction test for the dissection phenotype. We used the 2-week schedule previously reported, exposing the animals to the toxin prior to TGFβ signaling disruption at 11 weeks of age as decreasing ECM synthesis would impact the efficacy of BAPN (Figure 9A). Short-term BAPN administration did not affect blood pressure or induce hemorrhagic lesions in adult Tgfbr1/2^iSMCKO^ mice (Figure 9B,C). Histology was largely unremarkable though with a few focal breaks of elastic fibers similar to aortas from adult B6 WT mice exposed to BAPN (Supplemental Figure 9). In contrast, BAPN exposure for 14 days followed, a week later, by NE infusion for 7 days resulted in hypertension in Tgfbr1/2^iSMCKO^ mice (as did NE alone; cf. Figure 2B) with complete penetrance of the dissection phenotype, thus exceeding disease incidence with NE alone (Figure 9D,E). Hemorrhagic lesions were larger on gross examination with more extensive dissection by histology (Figure 9F,G).

**Figure 9:**
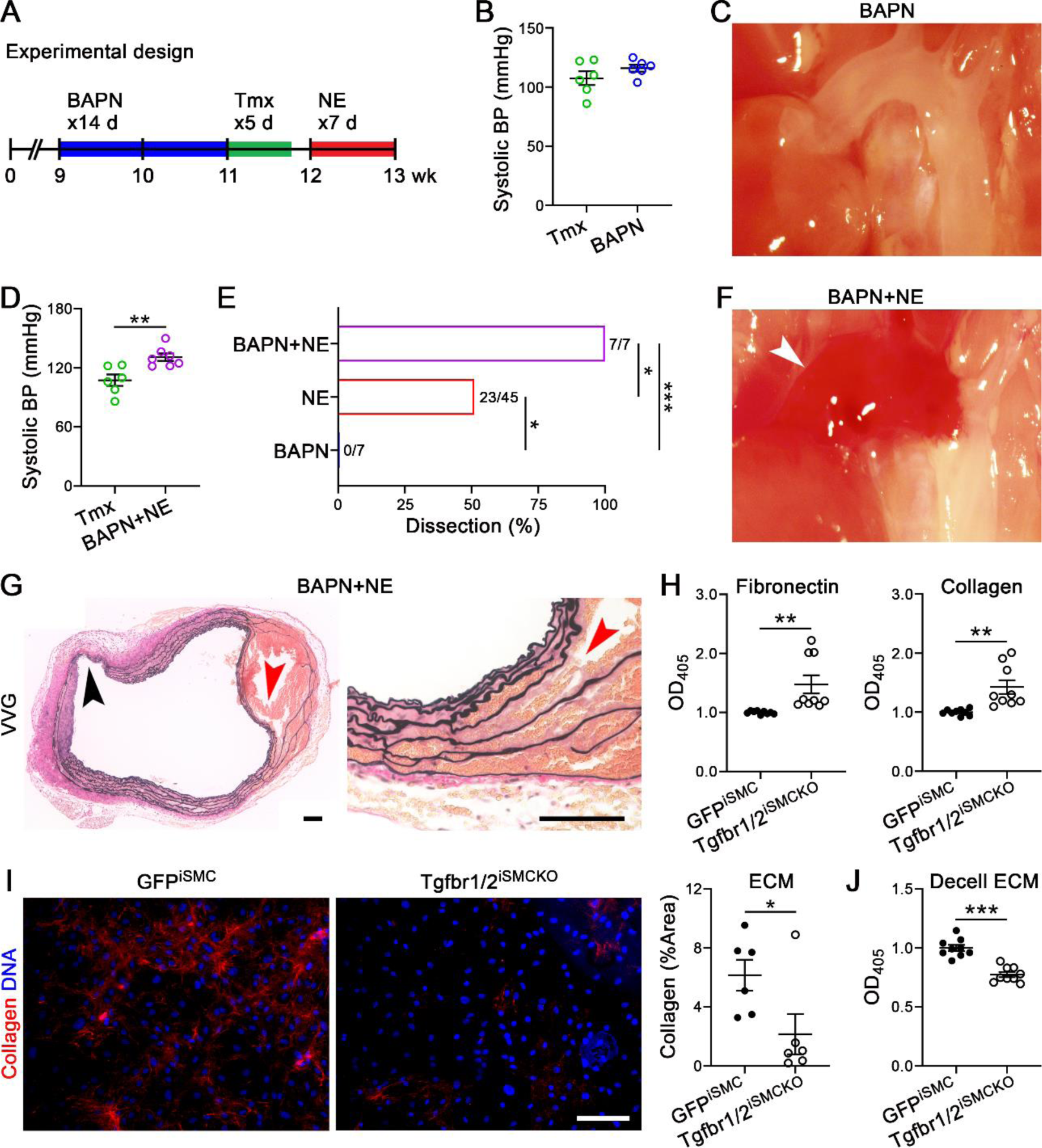
Impaired ECM synthesis predisposes to aortic dissection 1 week after disrupting TGFβ signaling in SMCs. (**A**) Nine-week-old Tgfbr1/2^iSMCKO^ mice were given BAPN at 150 mg/kg/d for 14 days, then tamoxifen (Tmx) for 5 days, without or with NE infusion at 3.88 μg/kg/min for 7 days, then examined at 12 or 13 weeks of age, respectively. (**B**) Systolic blood pressure (BP) measured by tail-cuff in Tgfbr1/2^iSMCKO^ mice given BAPN or not (*n* = 6). (**C**) In situ examination of BAPN-exposed animal showing thoracic aorta with unremarkable appearance. (**D**) Tail-cuff blood pressure in Tgfbr1/2^iSMCKO^ mice with BAPN+NE or not (*n* = 6-7). (**E**) Incidence of aortic dissection in BAPN- (0%, *n* = 7), NE- (51%, *n* = 45), and BAPN+NE- (100%, *n* = 7) infused Tgfbr1/2^iSMCKO^ mice. (**F**) In situ examination of BAPN+NE-infused animal showing large mural hematoma of ascending aorta and arch (white arrow). (**G**) Verhoeff–Van Gieson (VVG) stain confirming extensive dissection (red arrows) and intimomedial tear (black arrow) of ascending aorta, scale bars: 100 μm. (**H**) Colorimetric assay for number of GFP^iSMC^ and Tgfbr1/2^iSMCKO^ murine SMCs adherent to fibronectin- or collagen-coated plates after 1 h of plating (*n* = 9, pooled from 3 experiments); OD_405_ normalized to controls. (**I**) Confocal microscopy for CNA35-labeled collagen (red) and DAPI-labeled nuclei (blue) of SMC cultures from thoracic aortas of GFP^iSMC^ and Tgfbr1/2^iSMCKO^ mice, scale bar: 200 μm, and quantification of CNA35 labelling in culture ECM as %Area (*n* = 6). (**J**) Adhesion of Tgfbr1/2^iSMCKO^ murine SMCs to decellularized (Decell) ECM produced by cultured SMCs from GFP^iSMC^ and Tgfbr1/2^iSMCKO^ mice (*n* = 9, pooled from 3 experiments); OD_405_ normalized to controls. Data are shown as individual values with mean ± SEM, \**P* < 0.05, \*\**P* < 0.01, \*\*\**P* < 0.001 by t-test (B, D, H-J) or Fisher’s exact test between infusions (E).

Because these in vivo studies implicated ECM abnormalities as pathogenic, we investigated interactions of SMCs with ECM in vitro. Cultured aortic SMCs from Tgfbr1/2^iSMCKO^ mice displayed increased adhesion to exogenous fibronectin and collagen compared to control cells from GFP^iSMC^ mice (Figure 9H). Nevertheless, the mutant cells exhibited decreased adhesion to collagen-deficient, decellularized ECM elaborated by SMCs from Tgfbr1/2^iSMCKO^ mice versus control matrix from GFP^iSMC^ mice (Figure 9I,J). Together, these findings suggest that impaired adhesion of SMCs to defective ECM early after TGFβ signaling disruption sensitizes to pressure-induced aortic dissection.

## Discussion

In contrast to previous reports in mice that aortopathy presents with infusion of AngII but not NE (5,6), we found that the thoracic aorta dissected in adult Tgfbr1/2^iSMCKO^ mice with both AngII- and NE-induced elevations in blood pressure. Notwithstanding differences in mouse models (wild-type or germline hypercholesterolemic versus conditional disruption of TGFβ signaling), there are two critical differences between AngII and NE infusion. First, chronic delivery of AngII not only elevates blood pressure, it also stimulates a strong inflammatory response (25–27) that can compromise aortic wall properties (28); indeed, NE may even suppress an inflammatory response (29,30). Second, differential AngII type 1 receptor and α1-adrenoreceptor densities along the aorta (31–33) result in greater contractions of the thoracic aorta in response to phenylephrine (a NE analog) than to AngII. Hence, it is possible that thoracic aortic wall stress is greater in AngII-than NE-induced hypertension even at the same pressure (cf. Figure S5 in 34). In the present study, both chronic infusion and bolus delivery of AngII resulted in greater elevation of blood pressure than NE, despite selecting pressure-equivalent doses of vasoconstrictors based on previous work (5,6,19), which could have contributed to the higher rates of dissection with AngII. Although inflammatory effects appear to dominate in AngII-induced hypertension, with lesions exacerbated in cases of TGFβ neutralization (27), reduction of blood pressure with hydralazine nevertheless attenuates AngII-induced aortic inflammation, stiffening, and adventitial collagen deposition (35) and it prevented NE-induced dissection herein. Regardless of vasoactive agent, the role of pressure-elevated wall stress is undeniable; aortic damage occurs when mechanical stress exceeds material strength locally whether stress is normal and strength is compromised or stress is elevated and strength is normal, or both – which appears to be the case herein due to dissection triggered by high blood pressure following short-term loss of TGFβ signaling and associated changes in ECM transcripts.

We recognize several types of injury to the vessel wall that initiate and propagate dissection of the murine aorta: (i) intimomedial or entry tear, (ii) separation of SMCs from neighboring cells and fibrillar matrix by medial extravasation of blood, (iii) delamination (widening) of elastic laminae with traction on SMCs via intralaminar elastic fiber attachments, and (iv) further delamination of elastic laminae with fragmentation of SMCs and fibrillar matrix. Intimomedial tears extend through the intima and partially or fully through the media with the external elastic lamina and/or adventitia preventing transmural rupture. Notably, intimomedial tears can occur in isolation without extension of lesions to rupture or dissection along axial planes (36). RBCs infiltrate among SMCs rather than along elastic laminae suggesting medial vulnerability between cells and fibrillar matrix but not at cell attachments to intralaminar elastic fibers. Progressively, SMCs are stretched between widened elastic laminae changing their long axis from circumferential to radial and then tear with membrane fragments remaining attached to elastin. In areas of greatest delamination and RBC accumulation, the elastic laminae are often stripped clean of membrane and collagen fragments. The spectrum of abnormalities differs from predominant loss of SMC contacts to elastic fibers seen in aortopathy 10 weeks after conditional disruption of *Smad3* in SMCs of 6-week-old mice (37). Differences in medial injury in models of TGFβ signaling disruption may reflect a unique susceptibility of the developing versus mature aorta or a prolonged half-life of SMC-elastic fiber adhesion structures versus a shorter half-life of other SMC-ECM connections. Further temporal studies are necessary to characterize the turnover of diverse medial ECM proteins.

Although bulk passive mechanical properties of the mature ascending aortas were not statistically different between wild-type controls and Tgfbr1/2^iSMKO^ mice with 1 week of receptor disruption, nearly all trends (e.g., increases in wall thickness and decreases in both axial stretch and energy storage in Tgfbr1/2^iSMKO^ mice) were nevertheless consistent with statistically significant changes in descending thoracic aortas when TGFβ signaling was similarly disrupted at 11 weeks of age but evaluated mechanically after 3 weeks (23). Together, these results suggest a relatively slow but progressive deterioration of bulk properties with loss of TGFβ signaling in the adult mouse, consistent with a turnover of matrix that is characterized by decreased synthesis or increased degradation, or both. The former interpretation is supported by RNA sequencing, emphasizing that TGFβ is important not only for establishing the mechanical integrity of the developing aorta (10) but also for maintaining this integrity as part of normal homeostatic processes in maturity. Among the many genes that were downregulated, reductions in competent *Col3a1*, *Col5a2*, *Lox*, and *Bgn* are consistent with thoracic aortopathies in humans (38–40), with effects of some related pathologic variants confirmed in mice (41–44). Moreover, *COL15A1* variants associate with disease severity in families with syndromic thoracic aortic aneurysms, suggesting a genetic modifier role (45). The importance of these matrix constituents to structural integrity is thus clear.

The role of SMC contractility in this model is complex, however. We have previously reported decreased vasoconstrictive capacity of the aorta in our pressurized ex vivo vessel preparations consistent with decreased expression of contractile molecules several weeks after TGFβ signaling disruption (10,22,23). Others have reported opposing results of aortic hypercontractility (associated with endothelial dysfunction) and increased expression of contractile molecules several weeks after TGFβ signaling disruption in similar strains (11,46,47). Certain disparities may reflect inherent differences between our biaxial (isobaric, axially isometric) vessel and uniaxial ring tests (48), the latter which often do not account for differences in wall thickness and cannot account for altered axial stretch. In an unbiased time-series analysis of single-cell RNA sequencing of *Myh11* lineage-marked SMCs, loss of contractile phenotype was mild after 1 month, moderate at 2 months, and severe at 4 months following TGFβ signaling disruption in hypercholesterolemic mice (12). In the present study, we find minor changes in contractile molecules and contractility, besides regulatory enzymes such as *Mylk4*, within 1 week of receptor deletion, suggesting that delayed loss (or gain) in SMC contractile phenotype is not a direct consequence of TGFβ signaling alone. The mechanistic experiments to determine if changes in SMC contractility induce aortic vulnerability are limited by confounding effects on resistance vessels, except for Fasudil which did not affect blood pressure at the dose used and was the only inhibitor of myosin light chain activity to show a trend toward increased dissection. Pervasive effects of vasodilators on aortic and resistance arteriole SMCs (49) also confound the interpretation of previous work finding blood pressure-independent aortopathy in AngII-infused and *Nos3*-deficient mice (5,50). Furthermore, experimental designs over days to weeks with hydralazine may be susceptible to unrecognized transient high blood pressure and we did not undertake long-term vasodilator experiments.

Whereas most attention to wall integrity has focused on roles of elastic fibers and fibrillar collagens (I, III, V), the present data suggest that additional matrix and matricellular proteins may contribute substantially to wall integrity, including collagens IV, XV, and XVIII as well as CTGF. Types XV and XVIII collagen are multiplexins that, among other roles, associate with basement membranes and contribute to their integrity (51). Basement membrane proteins surround SMCs, albeit not uniformly, and are readily detected by immunofluorescence microscopy compared to the inconspicuous, electron-dense layer on electron microscopy (52,53). Impaired basement membrane constituents may facilitate SMC displacement by extravasated blood and thus compound defects in the fibrillar matrix. CTGF, a prototypical TGFβ-inducible product, directly induces ECM production and facilitates that by TGFβ (54); thus, reduction in CTGF may synergize with decreased transcription of ECM molecules after TGFβ signaling disruption. Further studies are required to elucidate the roles of diverse TGFβ-dependent ECM molecules in a vulnerable aorta phenotype and conditional deletion models can describe the kinetics of disease onset for proteins with varying turnover.

An essential role for TGFβ in the mature aorta is underappreciated and underscores the continuous turnover of ECM by vascular wall cells even under quiescent conditions after postnatal development completes by 8 weeks of age (17). Previous work reported only the absence of spontaneous aortic disease after receptor deletion in adult animals (10,55). The findings herein do not support mutually exclusive contractile versus synthetic SMC phenotypes but rather overlapping functions, including degradative properties required for protein turnover (56,57). The turnover of total collagen in the normal adult aortic wall, based on rodent studies with radiolabeled precursors, is slow with a half-life of 70 days that increases ∼4-fold with a reduction in half-life to 17 days with hypertension (58). Current methods to determine synthesis and degradation rates of individual proteins utilize nonradioactive, isotype-labeled amino acids and mass spectrometry, although less soluble ECM proteins are challenging to analyze. These techniques have not yet been applied to the vasculature, but protein half-lives in murine brain cortex ranged from 6, 25, 62, and 79 days for collagen XI, I, VI, and IV, respectively – other collagens were not identified (59). Besides differential turnover of individual collagens, a pool of newly synthesized collagen has more dynamic turnover than the persistent collagen network in tendons varying with the circadian cycle versus the lifetime of the organism, respectively (60). Though we did not assess protein degradation, the markedly decreased abundance of collagen XVIII within 7 days of TGFβ signaling disruption suggests that its normal balanced turnover is not preserved. Decreases in total collagen expression were modest (∼13%) 1 week after starting tamoxifen but in accordance with a greater reduction (∼30%) in the aortic media at 3 weeks after receptor deletion using similar histological analysis (23). While TGFβ is necessary for ECM production by SMCs at steady state, further studies are required to determine its role under dynamic conditions imposed by hypertensive remodeling, including probable effects on ECM degradation.

We conclude that animal models with pre-existent aortic vulnerabilities are more appropriate to test the clinical risk factors for aortic dissection of poorly controlled hypertension and extreme exertion. AngII or NE are relevant vasoconstrictors as both are elevated in neurohormonal response to exercise, resulting in increased blood pressure (61). The vulnerable aorta phenotype resulting from disruption of TGFβ signaling and sensitizing to dissection elicited by transient and sustained elevations in blood pressure provides a mechanism and experimental confirmation of recent computational modeling that hypertension exacerbates but does not initiate focal mural defects leading to aortopathy and that continual ECM turnover, either balanced or unbalanced, is required for progressive aortic disease (62). A central inference of our findings is that focal loss of medial collagen, rather than fibrosis that characterizes medial degeneration, associates with a vulnerable aorta phenotype. Conversely, accumulation of collagen within the media may afford some protection against dissection.

## Methods (analytical techniques are described in the Supplemental Materials)

### Mice

C57BL/6J mice (stock 000664) and *mT/mG* mice (stock 007676) were purchased from the Jackson Laboratory, *Myh11-CreER^T2^* mice were obtained from Dr. Stefan Offermanns, University of Heidelberg, *Tgfbr1*^f/f^ mice were obtained from Dr. Martin M. Matzuk, Baylor College of Medicine, and *Tgfbr2*^f/f^ mice were obtained from Dr. Harold L. Moses, Vanderbilt University. Compound mutant strains were bred to a B6 background for more than 6 generations (12,23). Male mice were euthanized at 12 or 13 weeks of age for analysis of thoracic aortas (the BAC containing *Myh11-CreER^T2^* inserts on the Y chromosome and female mice do not express the construct).

### Statistics

Graphs of quantitative data are generally presented as dot plots of individual values with overlying line and error bars representing the mean and SEM. Single numerical values are represented by columns. Comparison of continuous variables between 2 groups was by Student’s t-test, among more than 2 groups by 1-way ANOVA (and 1-way repeated measures ANOVA when the same subjects in each group) with Tukey’s multiple comparisons test for 1 independent variable, or 2-way ANOVA with Sidak’s multiple comparisons test for 2 independent variables, and of categorical variables between 2 groups was by Fisher’s exact test. Probability values were 2-tailed and *P* < 0.05 was considered to indicate statistical significance. Graph construction and statistical analyses were performed with Prism 9.2.0 (GraphPad Software).

### Study Approval

Research protocols were approved by the Institutional Animal Care and Use Committee of Yale University.

## Author Contributions

BJ, PR, CH, and GT designed the study. BJ, PR, CH, MW, S-IM, YC, GL, and LQ conducted experiments and acquired data. BJ, PR, CH, MW, S-IM, ABR, LQ, JDH, and GT analyzed and interpreted data. JDH and GT supervised the work. BJ, PR, RA, MAS, JDH, and GT wrote and edited the manuscript. The order of the three first authors was determined by relative effort: CH initiated the work, BJ continued the work, and PR completed the work, with all three collectively performing the bulk of the work and making critical contributions.

## Sources of Funding

This work was supported by grants from the NIH (R01 HL146723, R01 HL152187, R01 HL152197) and the Leducq Foundation (erAADicate Network).

## Notes

### Competing Interest Statement

The authors have declared no competing interest.

