## Supplemental_Materials for "Short-Term Disruption of TGFβ Signaling in Adult Mice Renders the Aorta Vulnerable to Hypertension-Induced Dissection"

### Expanded Methods

**Genotyping**. Ear DNA was isolated using DNeasy Blood & Tissue kits (#69506, QIAGEN). Genotyping was performed using the following PCR primers: *Myh11-CreER*$^{T2}$ (5'-TGA CCC CAT CTC TTC ACT CC-3', 5'-AAC TCC ACG ACC ACC TCA TC-3', and 5'-AGT CCC TCA CAT CCT CAG GTT-3'), *Tgfbr1*$^{f/f}$ (5'-ACT CAC ATG TTG GCT CTC ACT GTC-3' and 5'-AGT CAT AGA GCA TGT GTT AGA GTC-3'), *Tgfbr2*$^{f/f}$ (5'-TAA ACA AGG TCC GGA GCC CA-3' and 5'-ACT TCT GCA AGA GGT CCC CT-3'), *mT/mG* (5'-CTC TGC TGC CTC CTG GCT TCT-3', 5'-CGA GGC GGA TCA CAA GCAATA-3' and 5'-TCA ATG GGC GGG GGT CGT T-3').

**Infusions, Injections, and Treatments.** Cre-Lox recombination was induced by tamoxifen (T5648, Sigma-Aldrich) at 2 mg/d i.p. for 5 days starting at 11 weeks of age (or 1 mg/d i.p. for 5 days for a subgroup induced at 4 weeks of age), while Tgfbr1/2$^{+/+}$ controls were given a corn oil vehicle (C8267, Sigma-Aldrich). Some mice were infused with saline, NE (N5785, Sigma-Aldrich) at 3.88 μg/kg/min (1), or AngII (A9525, Sigma-Aldrich) at 1 μg/kg/min (1) s.c. for 7 days from 12 to 13 weeks of age; other mice were injected with single doses of saline, NE at 1.28 mg/kg (2), or AngII at 0.64 mg/kg (2) i.p. at 12 weeks of age. Additionally, some mice were treated with a single dose of hydralazine (1313006, Sigma-Aldrich) at 10 mg/kg (3) i.p. at 12 weeks of age, or treated with infusions of ML-7 (S8388, Selleckchem) at 73 μg/d (4), nifedipine (S1808, Selleckchem) at 5 mg/kg/d (5), fasudil (S1573, Selleckchem) at 50 mg/kg/d (6) s.c. for 7 days from 12 to 13 weeks of age, or infusion of BAPN (A3134, Sigma-Aldrich) at 150 mg/kg/d s.c. for 14 days from 9 to 11 weeks of age (7).

**In Situ Examination and Aorta Procurement.** Following anesthesia, with ketamine at 100 mg/kg and xylazine at 10 mg/kg i.p., and a midline incision, the abdomen and chest were opened widely, the circulation was flushed with saline via the left ventricle, then the mediastinum, lungs, and pulmonary artery were excised. The thoracic aorta was inspected for mural hematomas using a

SZX16 dissecting microscope with a camera attachment (Olympus). The thoracic aorta was excised from above the root to the diaphragm. In some cases, the descending thoracic aorta was cannulated and perfused retrograde with 10% formalin at 75 mmHg for 30 min after ligation of arch branches.

**Pump Implantation.** Osmotic minipumps (1007D for 7-day or 1002 for 14-day infusions, Alzet) were primed with drug overnight prior to implantation at 9 or 12 weeks of age. Under inhaled isoflurane anesthesia (1.5% at 1 L/min), an incision was made on the dorsum, the subcutaneous tissue was bluntly dissected, the pump implanted, and the incision closed with 6/0 non-absorbable suture. Buprenorphine at 0.1 mg/kg s.c. or i.p. was administered pre-operatively and q12h x 48h post-operatively. For animals that received two pumps (for BAPN and NE), the first pump was removed at 12 weeks of age when the second pump was implanted in a contralateral pocket.

**Blood Pressure.** Systolic pressure was measured noninvasively in conscious animals using a CODA volume-pressure recording sensor and occlusion tail-cuff (Kent Scientific Corporation). Mice were placed in warmed restraining chambers and pressures were recorded for 40 cycles after discarding the first 10 data points. Alternatively, the aorta was catheterized via the right carotid artery using a 1.0-Fr Mikro-Tip pressure catheter (Millar Instruments), and blood pressure was measured in lightly anesthetized mice using a PowerLab system (ADInstruments). After invasive pressure monitoring, the aortas were not used for microscopy studies due to possible injury.

**Ultrasound.** Transthoracic B-mode images of the ascending aorta were obtained in lightly isoflurane anesthetized animals using a Vevo 770 high-frequency ultrasound (VisualSonics). Luminal diameter at the level of the pulmonary artery was measured in triplicate at end-systole.

**Histology.** Thoracic aortas were fixed in 10% formalin at 4 °C overnight, transferred to 70% ethanol at 4 °C for 24-48 hours, and then segments were embedded in paraffin with the specimens oriented transversely. Five μm-thick sections were stained with hematoxylin and eosin, Verhoeff-Van Gieson, Masson's trichrome, Movat's pentachrome, or picrosirius red by Yale's Research Histology Laboratory using standard techniques. Medial collagen area fraction was quantified from picrosirius red birefringence in 4 color distributions using a custom MATLAB code (https://github.com/yale-cbl/histological-analysis).

**Confocal Microscopy.** Unpressurized or pressure-fixed ascending aortas were post-fixed in 10% formalin and embedded in paraffin or frozen in OCT and 5 μm-thick sections were incubated with antibodies to smooth muscle α-actin (FITC conjugate, F3777, Sigma-Aldrich), TER-119 (116202, BioLegend), integrin α8 (biotin conjugate, BAF4076, R&D), tdTomato-CNA35 (purified from plasmid #61606, Addgene), collagen I (72026, Cell Signaling), collagen I (Alexa Fluor 647 conjugate, 72827, Cell Signaling), collagen III (biotin conjugate, PA1-28532, Invitrogen), collagen IV (ab6586, Abcam), or perlecan (MA1-06821, Invitrogen) overnight at 4 °C. Unconjugated and biotinylated primary antibodies were detected with Alexa Fluor 405-, 568-, or 647-conjugated IgG or streptavidin (Invitrogen). Sections were covered with Alexa Fluor 633 hydrazide (A30634, Invitrogen) for 20 min (1:20,000) at room temperature to label elastin, then mounted with ProLong Gold Antifade reagent with DAPI (Life Technologies). Images were acquired with a SP8 confocal microscope and LAS X software (Leica).

**Western Blotting.** Thoracic aorta without hematomas were rapidly excised by sharp dissection to remove surrounding loose adipose tissue, then flash frozen. Protein was extracted from homogenized tissue using RIPA lysis buffer containing protease and phosphatase inhibitor cocktail tablets (Roche) and boiled in SDS sample buffer for 5 min. Equal amounts of protein per sample were separated by SDS-PAGE, transferred electrophoretically to a PVDF membrane (Bio-

Rad Laboratories), and blotted with antibodies to phospho-Smad2-S465/467 (3108, Cell Signaling), Smad2 (3103, Cell Signaling), phospho-MLC2-S19 (3675, Cell Signaling), MLC2 (3672, Cell Signaling), smooth muscle α-actin (ab5694, Abcam), and β-actin (A5316, Sigma-Aldrich) followed by horseradish peroxidase-conjugated secondary antibodies. Bound antibody was detected with Western Lightning Plus-ECL (Perkin Elmer). The blotting membrane was usually cut to allow the detection of multiple proteins of markedly different molecular weights. The appropriate molecular weight of each protein was confirmed at the time of analysis and was not annotated in the images.

**RT-PCR and Quantitative RT-PCR.** Aortas without hematomas were crushed, immersed in RLT lysis buffer (QIAGEN), and vigorously vortexed. Total RNA was isolated using a RNeasy Mini Kit and DNase Digestion Set (QIAGEN) according to the manufacturer's protocol. Reverse transcriptions were performed using an iScript cDNA Synthesis Kit (Bio-Rad). Quantitative RT-PCR was performed using a Bio-Rad CFX94 by mixing equal amount of cDNAs, Taqman gene master mix, and primers for *Mylk4* (Mm01161253_m1), *Mylk* (Mm00653039_m1), *Col15a1* (Mm00456551_m1), *Col18a1* (Mm00487131_m1), *Gapdh* (Mm99999915_g1), *Hprt* (Mm03024075_m1), and *Actb* (Mm02619580_g1) from Taqman. RNA-free ddH2O was used as negative controls for qPCR instead of cDNA samples. All reactions were in a 12.5 μL volume, in duplicate. PCR amplification consisted of 10 min of an initial denaturation step at 95 °C followed by 40 cycles of PCR at 95 °C for 15 s, and 60 °C for 1 min. We confirmed stable expression of several housekeeping genes.

**Flow Cytometry.** Thoracic aortas without hemorrhagic lesions were minced, incubated in 0.5 mL DMEM with 10% FBS, 1.5 mg/mL collagenase A (10103578001, Roche), and 0.5 mg/mL elastase (LS002294, Worthington Biochemical) 10 mg/mL *Bacillus licheniformis* protease (P5380, Sigma-Aldrich), 10 mg/mL Dispase II (D4693, Sigma-Aldrich), and 125 U/mL DNase I (DN25, Sigma-

Aldrich) for 3 h at 4 °C, and passed through a 40 μm filter (8). The cells were incubated with cell-impermeant viability dye (65-0865-14, Invitrogen) and blocked with unconjugated Fc receptor antibodies (139302,101302, and 149502, BioLegend) for 20 min at 4 °C. In some cases, the cells were fixed with Fixation Buffer (00-8222-49, eBioscience) for 15 min, washed, and resuspended in Permeabilization Buffer (00-8333-56, eBioscience). The cells were stained with FITC-anti-SMA (1:200, F3777, Sigma-Aldrich), biotinylated-anti-ITGA8 (1:200, BAF4076, R&D), calnexin (1:200, ab22595, Abcam) or irrelevant IgG as controls for 30 min on ice in the dark. Unconjugated and biotinylated antibodies were labeled with Alexa Fluor 405-conjugated IgG (A48258, Invitrogen) or Alexa Fluor 647 conjugated streptavidin (S21374, Invitrogen). The cells were pelleted and resuspended in 0.4% BSA/PBS for analysis using a LSR II (BD Biosciences) and the data was analyzed with FlowJo software.

**Transmission Electron Microscopy.** Ascending aortas of 12-week-old Tgfbr1/2$^{iSMCKO}$ mice, with and without dissection, were perfusion-fixed with 10% formalin at 75 mmHg for 30 min after ligation of arch branches and then post-fixed in 2% PFA and 2.5% glutaraldehyde in 0.1 M sodium cacodylate buffer at pH 7.4 for 1 h at room temperature, then stored in the same buffer at 4 °C until processing. Specimens were post-fixed in 1% $OsO_4$ in the same buffer at room temperature for 1 h. After en bloc staining with 2% aqueous uranyl acetate for 1 h, the tissue was dehydrated in a graded series of ethanol to 100%, followed by propylene oxide, then embedded in EMbed 812 resin, and the sample blocks were polymerized at 60 °C overnight. Thin sections (60 nm) were cut with an EM UC7 ultramicrotome (Leica) and post-stained with 2% uranyl acetate and lead citrate. Sample grids were examined in a Tecnai G2 Spirit BioTwin transmission electron microscope (FEI) at 80 kV of accelerating voltage and digital images were recorded with a SIS Morada CCD camera (Olympus) with iTEM imaging software.

**Biomechanical Assessment.** A biaxial testing device was used as described (9,10). In brief, excised ascending aortas, with and without dissection, were cleaned of perivascular tissue, cannulated on custom-drawn glass pipets, secured with sutures at each end, mounted on a biaxial testing device, and submerged in Krebs-Ringer's solution at 37 °C oxygenated with 95% $O_2$ / 5% $CO_2$ to maintain a physiologic pH.

*Active Mechanical Contraction Testing:* The viability of the active tone in the aorta was assessed through two initial contractions to 100 mM KCl at two different combinations of pressures and vessel lengths followed by washouts with normal Krebs-Ringer solution. The vessels were then set to 90 mmHg and the specimen-specific value of in vivo axial stretch and contracted with 100 mM KCl for 15 min followed by 10 min of relaxation by washout. This was repeated for 1 µM phenylephrine.

*Passive Mechanical Biaxial Testing*: After completing the active testing, the testing chamber was drained and refilled with Hanks' buffered salt solution and maintained at room temperature to minimize smooth muscle contractility. Vessels were mechanically preconditioned, by cyclic pressurization between 10 to 140 mmHg at the estimated in vivo value of axial stretch, to minimize viscoelastic contributions to the mechanical behavior. The aortic segments were then subjected to a series of seven biaxial protocols consisting of cyclic pressurization from 10 to 140 mmHg while the vessel was held fixed at three different axial stretches (95, 100, and 105% of the in vivo value), and cyclic axial stretching at four fixed pressures (10, 60, 100, and 140 mmHg).

*Data Analysis of Active and Passive Mechanical Properties*: The contractile properties were assessed by calculating changes in inner radius and mean circumferential stress between relaxed and contracted states. The passive pressure-diameter and axial force-length data were fit with a validated four-fiber family constitutive model via a nonlinear regression (Levenberg-Marquardt) of a data set from all seven testing protocols. Specifically, we used a Holzapfel-type nonlinear stored energy function $W$,

$$W(\mathbf{C}, \mathbf{M}^i) = \frac{c}{2}(I_{\mathbf{C}} - 3) + \sum_{i=1}^{4} \frac{c_1^i}{4c_2^i}\left\{\exp\left[c_2^i\left(IV_{\mathbf{C}}^i - 1\right)^2\right] - 1\right\}, \qquad (xx)$$

where $c$, $c_1^i$, and $c_2^i$ ($i = 1,2,3,4$ denote the four predominant fiber family directions) are material parameters, with $c$ and $c_1^i$ having units of stress (kPa) and $c_2^i$ dimensionless. $I_{\mathbf{C}} = tr(\mathbf{C})$ and $IV_{\mathbf{C}}^i = \mathbf{M}^i \cdot \mathbf{C}\mathbf{M}^i$ are coordinate invariant measures of the finite deformation, computed in terms of the right Cauchy-Green tensor $\mathbf{C} = \mathbf{F}^{\mathrm{T}}\mathbf{F}$ where the deformation gradient tensor $\mathbf{F} = \mathrm{diag}[\lambda_r, \lambda_\theta, \lambda_z]$, with $det\mathbf{F} = 1$ because of assumed incompressibility. The direction of the $i^{th}$ family of fibers is identified by the vector $\mathbf{M}^i = [0,\ \sin\alpha_0^i,\ \cos\alpha_0^i]$, with the model parameter $\alpha_0^i$ denoting a fiber angle relative to the axial direction in the traction-free reference configuration. Values of biaxial stress and material stiffness were computed from appropriate differentiation of the stored energy function.

**Bulk RNA-Seq.** Total RNA was isolated from *Myh11* lineage-marked SMCs or thoracic aortas without hemorrhagic lesions as described for RT-PCR and quality control was assessed by nanodrop and an Agilent Bioanalyzer. Next-generation, whole-transcriptome sequencing was performed using a NovaSeq 6000 System (Illumina) at the Yale Center for Genome Analysis. Low-quality reads were trimmed, and adaptor contamination were removed using Trim Galore (v0.5.0). Trimmed reads were mapped to the mouse reference genome (GRCm38) using HISAT2 (v2.1.0) (11). Gene expression levels were quantified using StringTie (v1.3.3b) (12) with gene models (M15) from the GENCODE project. Differentially expressed genes were identified using DESeq2 (v 1.22.1) (13). The raw data for SMCs (Figure 6) and aortas (Supplemental Figure 6) is deposited in Gene Expression Omnibus (GEO) GSE194085 with processed data in appended Excel tables.

**Single-Cell RNA-Seq.** Cells were isolated from the thoracic aorta without hemorrhagic lesions as described for flow cytometry and incubated with cell-impermeant viability dye (Thermo Fisher) for

20 min at 4 °C. Viable, GFP+ SMCs were sorted with a FACSAria (BD Biosciences) and collected in 0.4% BSA/PBS. The selected cells were processed for single-cell RNA-seq library preparation using the Chromium™ Single Cell Platform (10x Genomics) as per the manufacturer's protocol. Briefly, single cells were partitioned into Gel Beads in Emulsion using the Chromium™ system (10x Genomics), followed by cell lysis and barcoded reverse transcription of RNA, cDNA amplification and shearing, and 5' adaptor and sample index attachment. Single-cell RNA-seq libraries were sequenced on a NovaSeq 6000 System (Illumina) at the Yale Center for Genome Analysis. The data was processed to obtain fastq sequences using the "cellranger mkfastq" program provided by the vendor (10x Genomics). The reference mouse genome (mm10) was customized by addition of gene annotations for the exogenous tdTomato and eGFP genes. Raw reads were aligned to the customized mouse reference genome and gene expression was quantified using the "cellranger count" program provided by the vendor (10x Genomics). Gene expression data was further processed using the Seurat package (version 4.0), including data filtering, normalization, data dimension reduction, cell clustering, marker identification, cell type identity assignment, and data visualization. Filtering criteria for genes were: (i) include genes detected in at least 10 cells. Filtering criteria (satisfy all criteria) for cells were: (i) include cells with eGFP > 1, (ii) include cells > 200 genes detected, (iii) include cells with transcripts between 5,000 and 25,000, and (iv) include cells with mitochondrial RNA < 10%. The data was normalized using SCTransform (https://satijalab.org/seurat/articles/sctransform_vignette.html). Differential gene expression was used for clustering by T-distributed Stochastic Neighbor Embedding (tSNE) or Uniform Manifold Approximation and Projection (UMAP) and these projections were used for visualization with dimensional reduction. The raw data for SMCs (Figure 6) and aorta cells (Supplemental Figure 6) is deposited in GEO GSE194085 with processed data in appended Excel tables.

**Gene Ontology Enrichment Analysis.** Differentially expressed genes between experimental groups with log2(fold change) > 1 and false discovery rate-adjusted $P \leq 0.05$ were used for gene ontology enrichment analysis of bulk and single-cell RNA-seq datasets. Upregulated and downregulated genes were separately analyzed using DAVID v6.8 (https://david.ncifcrf.gov/) to identify enriched biological themes among biological process, cellular component, and molecular function terms. Enriched terms were ranked by *P*-value and the top 10 in each category displayed.

**Cell Culture.** Thoracic aortas without hemorrhagic lesions were digested for 5 min at 37 °C in HBSS containing 1 mg/mL collagenase A (10103578001, Roche) to promote sharp removal of the adventitia under a dissecting microscope. The denuded vessels were transferred into 0.5 mL DMEM (Thermo Fisher) containing 10% FBS (Life Technologies), 1.5 mg/mL collagenase A, and 0.5 mg/mL elastase (LS002294, Worthington Biochemicals) and incubated at 37 °C for 30 min while the digest was triturated with a pipette every 15 min. The mixture was centrifuged, then the cells were resuspended in Claycomb medium (Sigma-Aldrich) supplemented with 10% FBS, and cultured in 35 mm dishes in a $CO_2$ incubator at 37 °C. On reaching confluence, the cells were dissociated with trypsin/EDTA and GFP+ SMCs were sorted under sterile conditions using a FACSAria (BD Biosciences). The selected cells were expanded in DMEM with 10% FBS and used for experiments at passage 1 to 3. In certain experiments, SMCs were plated onto cover slips in 12-well plates, cultured for 4 d, washed twice, fixed with 4% paraformaldehyde, and collagen was labeled with the CNA35 probe overnight at 4 °C. The slides were incubated with Pro-Long Gold Mounting Reagent with DAPI (Life Technologies) and immunofluorescence images were acquired using an Axiovert 200M microscopy system (Carl Zeiss MicroImaging).

**Decellularized ECM.** Previously described techniques for endothelial cells and fibroblasts were adapted for SMCs (14,15). In brief, early passage SMCs were plated on uncoated plastic and expanded to confluence. The medium was not changed and acquired an acidic pH color within 7-

10 days. The cells were lysed with PBS containing 0.5% Triton X-100 and 20 mM $NH_4OH$ at room temperature for 10 min and the plates were washed 3 times with PBS. Decellularization was confirmed by microscopy. Decellularized matrices were stored at 4 °C for up to 2 weeks until use.

**Adhesion Assays.** Ninety-six well flat-bottom plates (351172, Falcon) were coated with purified bovine fibronectin (150025, MP Biomedicals) or rat-tail collagen (354249, Corning) at 6 µg/mL in 50 µL PBS at 4 °C overnight. Alternatively, decellularized matrices within the 96 well flat-bottom plates in which they were formed were used instead of exogenous fibronectin or collagen. The plates were washed once with PBS, blocked with freshly prepared, heat denatured BSA (A9418, Sigma-Aldrich) at 10 mg/mL dissolved in PBS for 1 h, and washed twice. Early passage SMCs were trypsinized, the enzymatic activity stopped with soybean trypsin inhibitor, and resuspended in serum-free DMEM containing 1 mg/mL BSA. The cells were seeded at $3 \times 10^4$ cells in 100 µL solution per well, incubated for 1 h at 37 °C, and washed twice with PBS containing 1 mM $Ca^{2+}$ and 2 mM $Mg^{2+}$. The number of adherent cells was determined by a colorimetric assay in which 100 µL of nitrophenyl phosphate (P4744, Sigma-Aldrich) at 3 mg/mL in 50 mM sodium acetate, pH 5.0 plus 0.4% Triton X-100 was added to each well, incubated at room temperature for 1 h, then 50 µL NaOH at 1M was added to each well and the optical density (OD) was determined spectrophotometrically at 405 nm. OD values were corrected for empty well readings and when pooled from several experiments, the results were normalized to wild-type controls because of batch variability.

**Protein Purification.** The plasmid pET28a-tdTomato-CNA35 encoding tdTomato-CNA35 was purchased from Addgene (61606) and purified as described previously (16). The plasmid was transformed into BL21 competent cells (EC0114, Thermo Fisher). Single colonies were picked to inoculate 6 mL LB medium containing 10 g/L peptone (211677, Thermo Fisher), 10 g/L NaCl, 5 g/L yeast extract (AB01208-02000, AmericanBio) supplemented with 25 µg/mL kanamycin

(AB01100-00010, AmericanBio). The bacteria were grown overnight at 37 °C then transferred and grown in 300 mL LB medium containing 25 µg/mL kanamycin. When the optical density value at 600 nm reached ~0.6, 0.2 mM IPTG (I6758, Sigma-Aldrich) was added and incubated for ~12 hours at 30 °C. The cultures were centrifuged, and the pellets resuspended in 10 mL Bugbuster (70584, Millipore) containing 10 µL benzonase (70750, Millipore) and 5 mM imidazole (I2399, Sigma-Aldrich) and incubated for 30 min at room temperature. The suspension was centrifuged and the protein in the supernatant was purified by Econo-Pac chromatography columns (7321010, Bio-Rad). Three milliliter Ni-NTA agarose resin (70666, Millipore) was loaded to the column and subsequently washed with 2 column volumes of wash buffer I (20 mM Tris-HCl pH 7.9, 0.5 M NaCl, and 30 mM imidazole). The supernatant was loaded onto the column and washed with 4 column volumes of wash buffer II (20 mM Tris-HCl pH 7.9, 0.5 M NaCl, and 60 mM imidazole). The protein was eluted by 4 column volumes of elution buffer (20 mM Tris-HCl pH 7.9, 0.5 M NaCl, and 500 mM imidazole). The buffer in which the protein was dissolved was changed to 50 mM Tris-HCl pH 8.0 and 100 mM NaCl by repeated concentration and dilution steps using Amicon Ultra-4 Centrifugal Filter Units (UFC801008, Millipore). The protein was kept in the dark at 37 °C for 16 h to allow complete chromophore maturation.

**Expanded Methods References**

14. Vlodavsky I. Preparation of extracellular matrices produced by cultured corneal endothelial and PF-HR9 endodermal cells. *Curr Protoc Cell Biol*. 2001;Chapter 10:Unit 10.4.

15. Franco-Barraza J, Beacham DA, Amatangelo MD, Cukierman E. Preparation of Extracellular Matrices Produced by Cultured and Primary Fibroblasts. *Curr Protoc Cell Biol*. 2016;71:10.9.1-10.9.34.

16. Aper SJ, van Spreeuwel AC, van Turnhout MC, van der Linden AJ, Pieters PA, van der Zon NL, de la Rambelje SL, Bouten CV, Merkx M. Colorful protein-based fluorescent probes for collagen imaging. *PLoS One*. 2014;9(12):e114983.

### Supplemental Figures

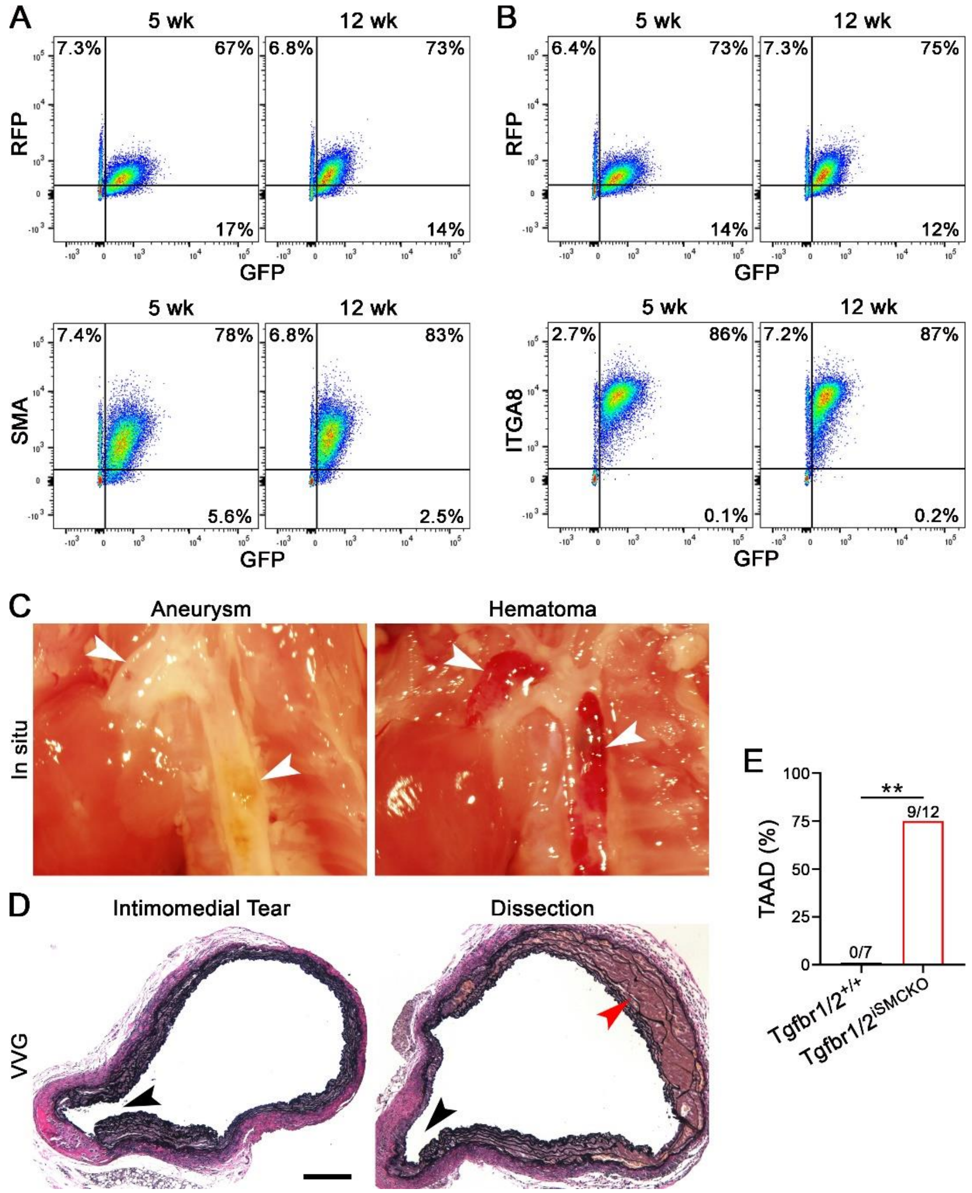


**Supplemental Figure 1: Disruption of TGFβ signaling in SMCs of immature aortas.** Cells were enzymatically isolated from thoracic aortas of 5- and 12-week-old Tgfbr1/2$^{iSMCKO}$ mice induced with tamoxifen over 5 days starting at 4 and 11 weeks of age, respectively, and analyzed by flow cytometry. (**A**) Red fluorescent protein (RFP), GFP, and smooth muscle α-actin (SMA) expression in fixed, permeabilized cells. (**B**) RFP, GFP, and integrin α8 (ITGA8) expression in non-permeabilized cells. SMCs with Cre-mediated recombination express both GFP and cell type-specific markers (SMA or ITGA8) and represent > 90% of total SMCs in both immature and mature aortas; GFP+ SMCs also express RFP, but at lower levels than cells without recombination, because of persistent protein expression 2-7 days after tamoxifen (RFP is absent several weeks after induction). Alternatively, 4-week-old *Tgfbr1*$^{f/f}$.*Tgfbr2*$^{f/f}$.*Myh11-CreER*$^{T2}$.*mT/mG* mice were injected with vehicle (Tgfbr1/2$^{+/+}$) or tamoxifen (Tgfbr1/2$^{iSMCKO}$) for 5 days and ascending aortas were examined at 8 weeks of age. (**C**) In situ examination showing aortic aneurysms and mural hematomas (white arrows) in Tgfbr1/2$^{iSMCKO}$ mice. (**D**) Verhoef-Van Gieson (VVG) stains of ascending aortas revealing intimomedial tears (black arrows) and dissection (red arrow) in Tgfbr1/2$^{iSMCKO}$ mice, scale bar: 200 µm. (**E**) Incidence of thoracic aorta aneurysm and dissection (TAAD) in 8-week-old Tgfbr1/2$^{+/+}$ ($n$ = 0/7) and Tgfbr1/2$^{iSMCKO}$ ($n$ = 9/12) mice without vasoconstrictor administration, $^{**}P < 0.01$ by Fisher's exact test.

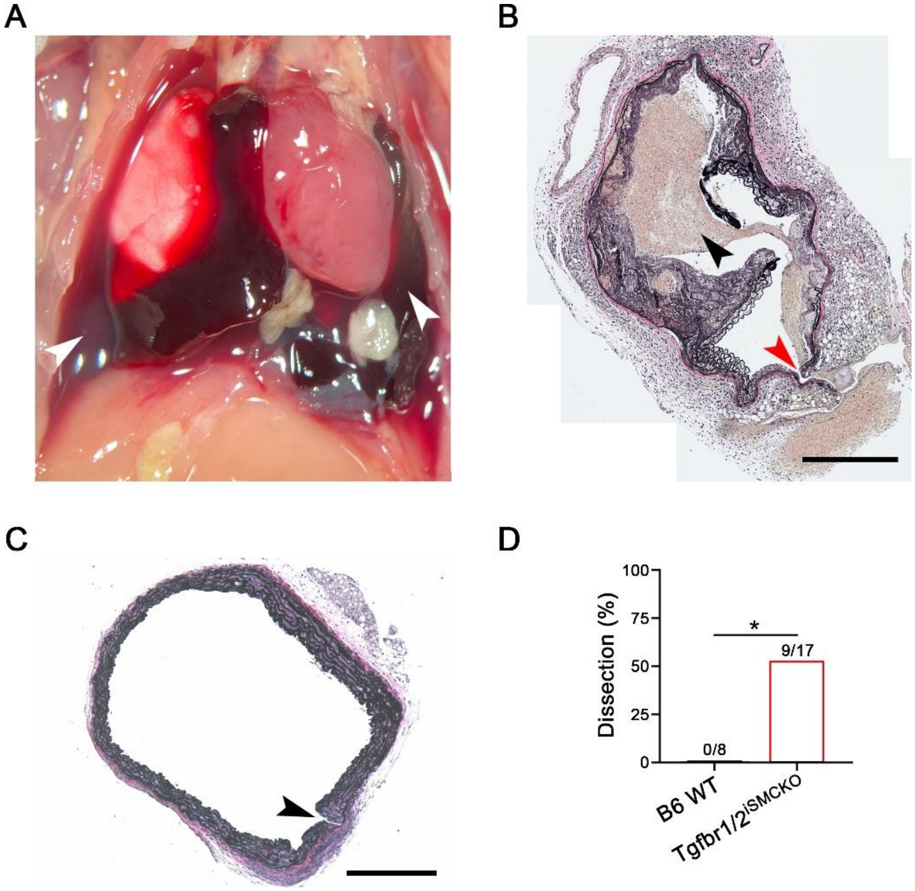


**Supplemental Figure 2: Spectrum of vasoconstrictor-induced aortic tears.** Twelve-week-old mice were infused with NE at 3.88 µg/kg/min or AngII at 1 µg/kg/min s.c. by osmotic minipump for 7 days. (**A**) Post-mortem examination after sudden death of Tgfbr1/2$^{iSMCKO}$ mouse at 6 days of AngII infusion revealing hemothorax (white arrows). (**B**) Verhoeff-Van Gieson stain of descending thoracic aorta of this animal showing contained rupture with false lumen (black arrow) and channel of free rupture connecting the lumen to extravascular blood accumulation (red arrow), merged image of multiple high magnification photomicrographs, scale bar: 1 mm. (**C**) Verhoeff-Van Gieson stain of ascending thoracic aorta of NE-infused Tgfbr1/2$^{iSMCKO}$ mouse without visible hematoma showing a discrete intimomedial tear (black arrow) but no evidence of blood extravasation between elastic lamellae, scale bar: 300 µm. (**D**) Incidence of aortic dissection in B6 WT (*n* = 0/8) and Tgfbr1/2$^{iSMCKO}$ (*n* = 9/17) mice after NE infusion for 1 week; *$P$ < 0.05 by Fisher's exact test.

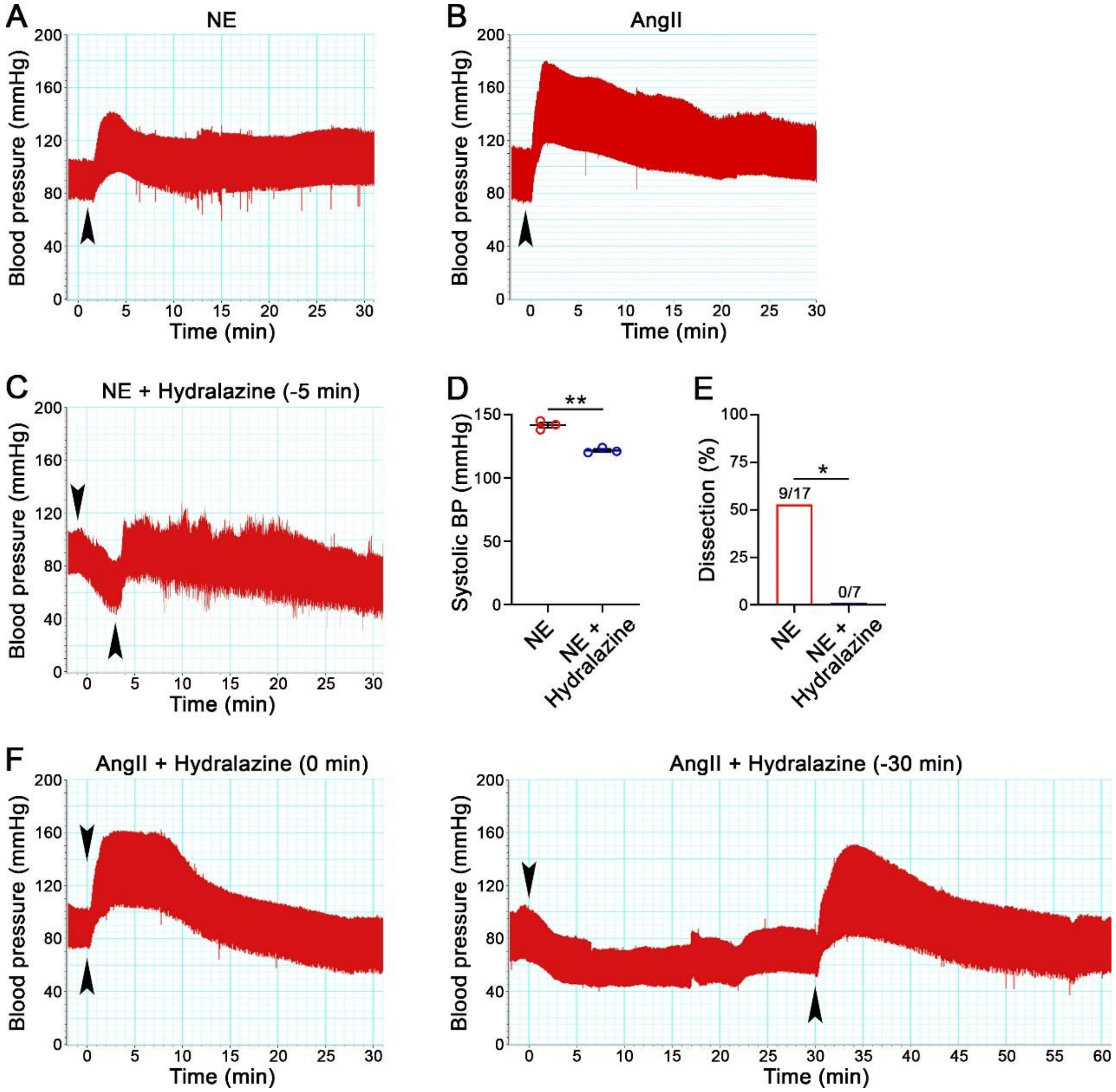


**Supplemental Figure 3: Hydralazine prevents NE-mediated pressure elevation and aortic dissection.** Millar catheter measurement of central blood pressure (BP) after intraperitoneal injection of (**A**) NE at 1.28 mg/kg or (**B**) AngII at 0.64 mg/kg (arrows), both monitored thereafter for 30 minutes. (**C**) Similar assessment of blood pressure after intraperitoneal injection of hydralazine at 10 mg/kg 5 minutes prior to administration of NE (arrows). (**D**) Maximum systolic BP over 30 minutes after injection ($n$ = 3). (**E**) Incidence of aortic dissection in 12-week-old Tgfbr1/2$^{iSMCKO}$ mice injected with NE ($n$ = 9/17) or NE plus hydralazine ($n$ = 0/7) after 30 minutes. (**F**) Blood pressure after intraperitoneal injection of hydralazine at 10 mg/kg from 0 to 30 minutes prior to administration of AngII (arrows); varying pretreatment protocols were ineffective at the doses tested unlike the successful prevention of NE-induced hypertension. Data are shown as individual values with mean ± SEM; *$P$ < 0.05, **$P$ < 0.01 by t-test (panel D) or Fisher's exact test (panel E).

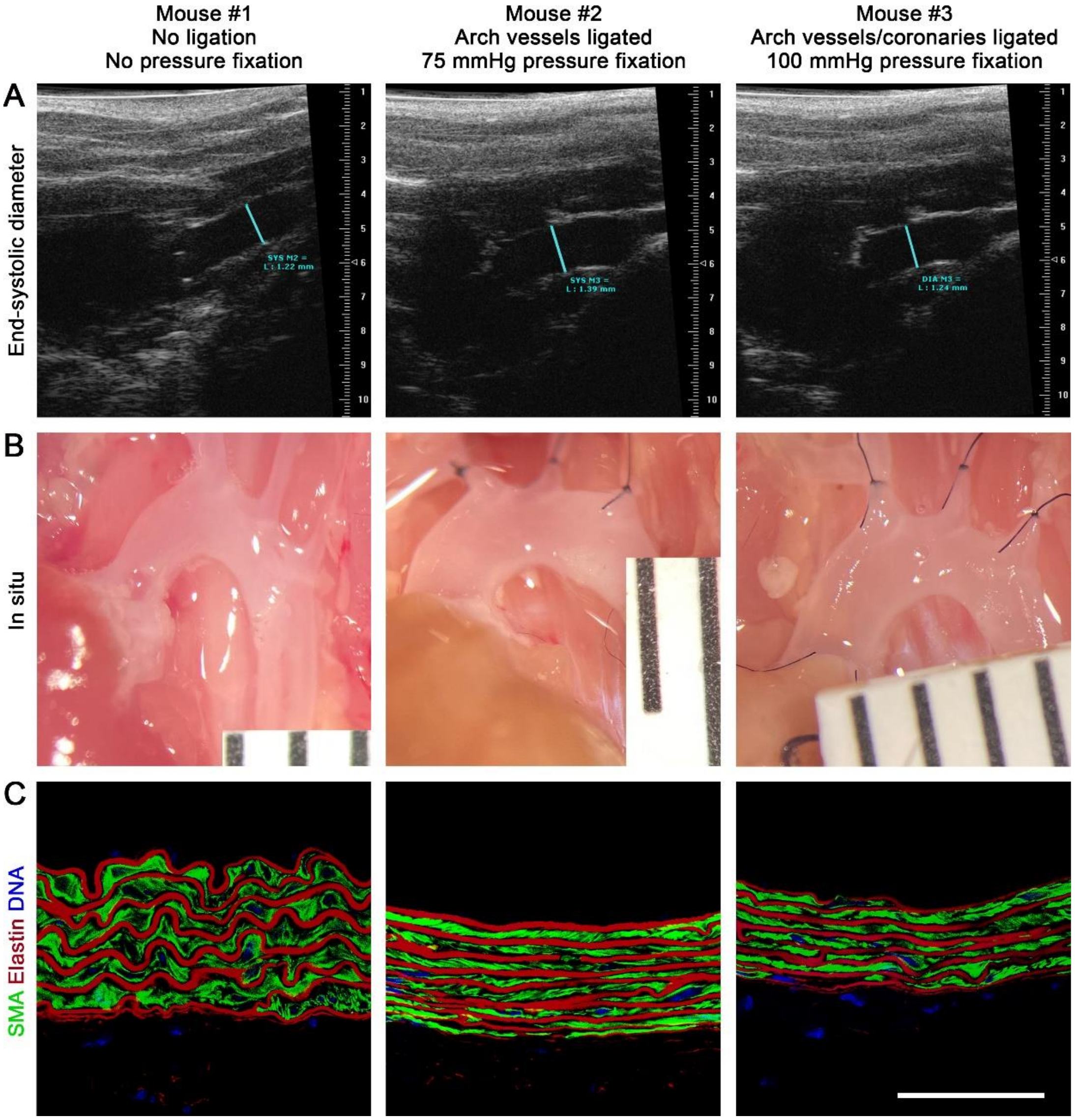


**Supplemental Figure 4: Pressure fixation of murine thoracic aortas.** (**A**) B6 WT mice of various ages were examined by ultrasound to determine end-systolic diameter (blue line) of the ascending aorta; ruler calibration in mm. (**B**) The circulation was flushed with saline via the left ventricle and the thoracic aorta remained unpressurized (left panel) or the descending thoracic aorta was cannulated and perfused with 10% formalin for 30 minutes at room temperature after ligation of the arch vessels (middle panel) or both the arch vessels and coronary arteries (right panel). The perfusion solution was at a height of 136 cm (equivalent to 100 mmHg) and the perfusion pressure was 75 mmHg after ligation of the arch vessels alone (likely losing pressure via runoff in the coronary circulation) and 100 mmHg after ligation of both arch vessels and coronary arteries. The in situ ascending aorta diameter after the procedure as a fraction of end-systolic diameter was 89% with no perfusion fixation, 98% with arch vessels ligated and perfusion pressure of 75 mmHg, and 105% with both arch vessels and coronary arteries ligated and perfusion pressure of 100 mmHg; ruler markings: 1 mm. (**C**) Confocal microscopy performed after labelling of SMCs with smooth muscle α-actin (SMA) antibody, elastin with AF633 hydrazide, and nuclei with DAPI revealed straightening of the medial laminae after pressure fixation, scale bar: 50 µm. The technique of arch vessel ligation with perfusion pressure of 75 mmHg was selected for further experiments to avoid overdistension of aortas in situ.

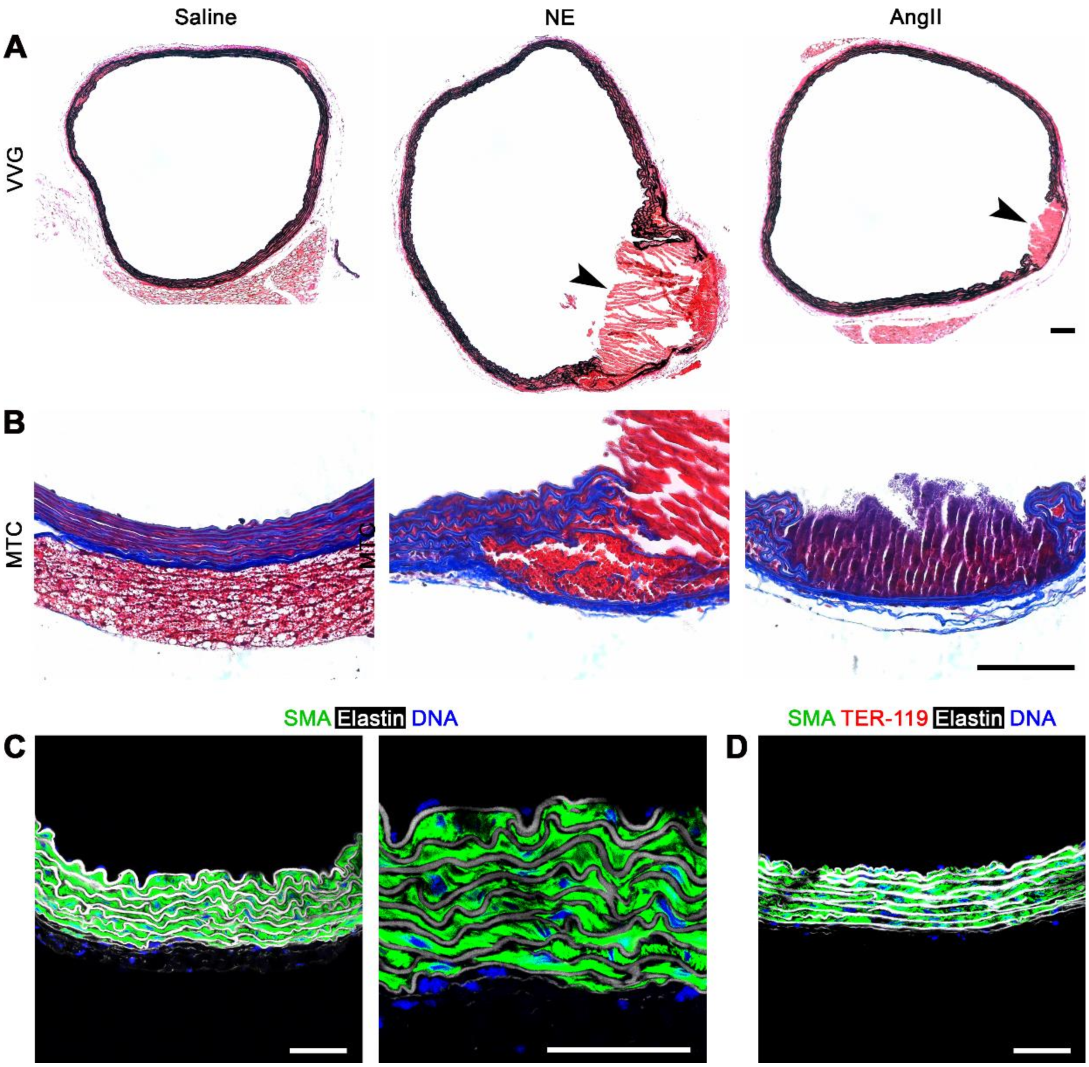


**Supplemental Figure 5: Histology of dissected and non-dissected aortas.** Microscopic structure of ascending aortas was examined following TGFβ signaling disruption at 11 weeks of age and i.p. injection of 12-week-old Tgfbr1/2$^{iSMCKO}$ mice with saline, NE, or AngII for 30 minutes. (**A**) Verhoeff-Van Gieson (VVG) stains show fresh thrombus (arrows) plugging entry tears that extend to or through the external elastic lamina. (**B**) Masson's trichrome (MTC) stains show disruption of medial collagen fibers (blue color) together with adjacent elastic fibers (black color in VVG stain), but intact adventitial collagen fibers preventing free rupture. (**C**) Confocal microscopy after labeling smooth muscle α-actin (SMA) for SMC cytoskeleton, AF633 hydrazide for elastin, and DAPI for nuclei shows orderly SMCs attached to elastic laminae after saline injection. (**D**) Additionally, TER-119 labelling for RBCs shows absence of dissection in NE-treated aortas without hemorrhagic lesions. Pressure-fixed (A, B, D) and unpressurized (C) specimens. Scale bars: 100 μm (A, B) and 50 μm (C, D).

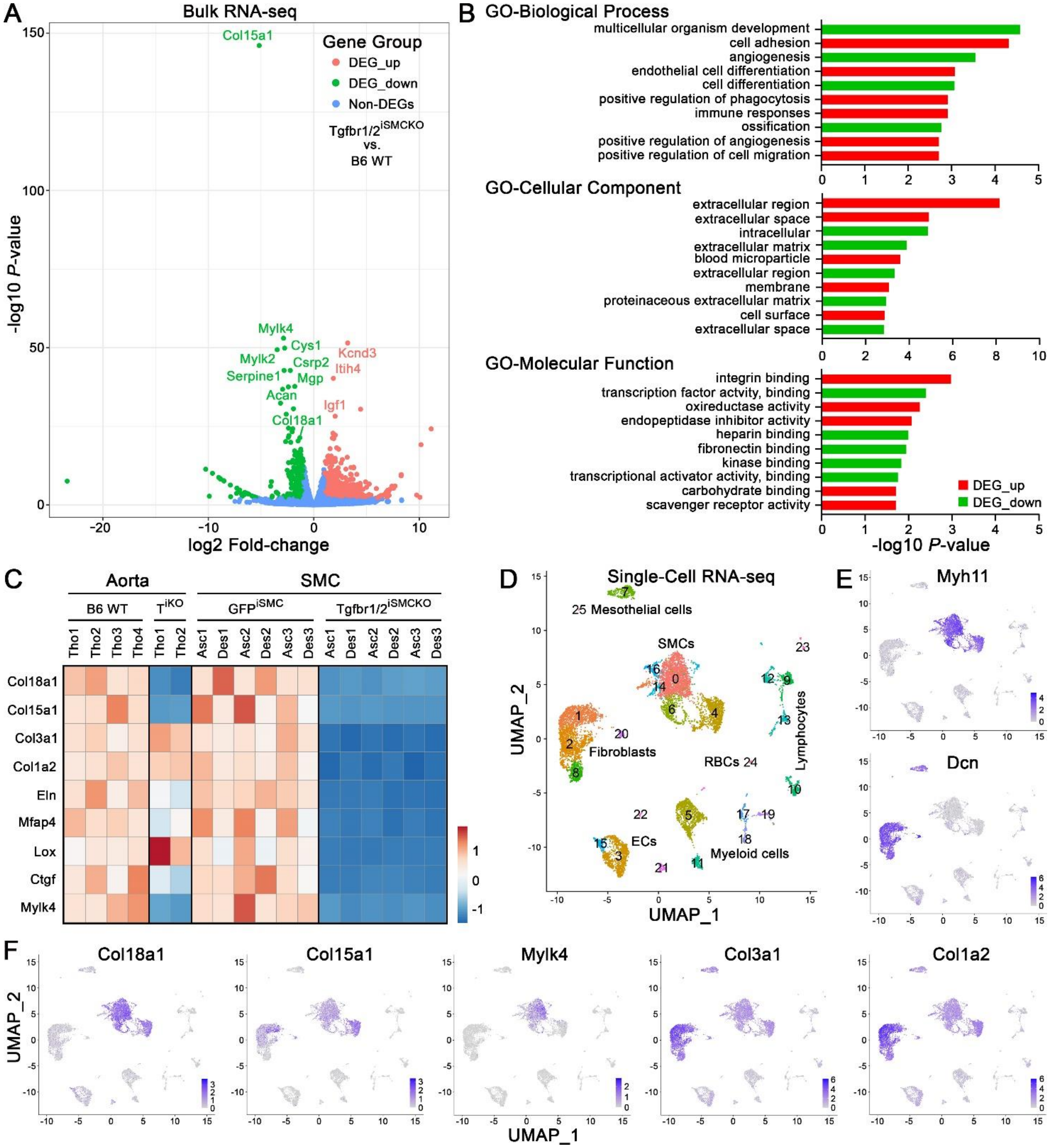


**Supplemental Figure 6: Transcriptome profiling of whole aortas versus isolated SMCs.** Bulk RNA-seq of thoracic aortas from 12-week-old B6 WT mice (*n* = 4) and Tgfbr1/2$^{iSMCKO}$ mice (*n* = 2) 1 week after TGFβ receptor disruption shown as (**A**) volcano plot highlighting differentially expressed genes (DEG) and (**B**) gene ontology (GO) enrichment analysis. (**C**) Heatmap comparing ECM and regulatory contractile molecules by bulk RNA-seq of thoracic aortas (Tho, *n* = 6) versus SMCs isolated from ascending/arch (Asc, *n* = 6) and descending (Des, *n* = 6) segments from 12-week-old GFP$^{iSMC}$ and Tgfbr1/2$^{iSMCKO}$ (T$^{iKO}$) mice illustrating common regulation of only limited genes. (**D**) Single cell RNA-seq of aortic cells from 12-week-old GFP$^{iSMC}$ mice (*n* = 2) with UMAP plot differentiating 25 clusters, including SMCs and fibroblasts identified by (**E**) cell type-specific markers *Myh11* and *Dcn*, respectively. (**F**) *Col18a1*, *Col15a1*, and *Mylk4* (differentially expressed by bulk RNA-seq of both aortas and SMCs) are preferentially expressed in SMCs, whereas *Col3a1* and *Col1a2* (differentially expressed by bulk RNA-seq of SMCs but not aortas) predominate in fibroblasts thus confounding whole tissue analysis.

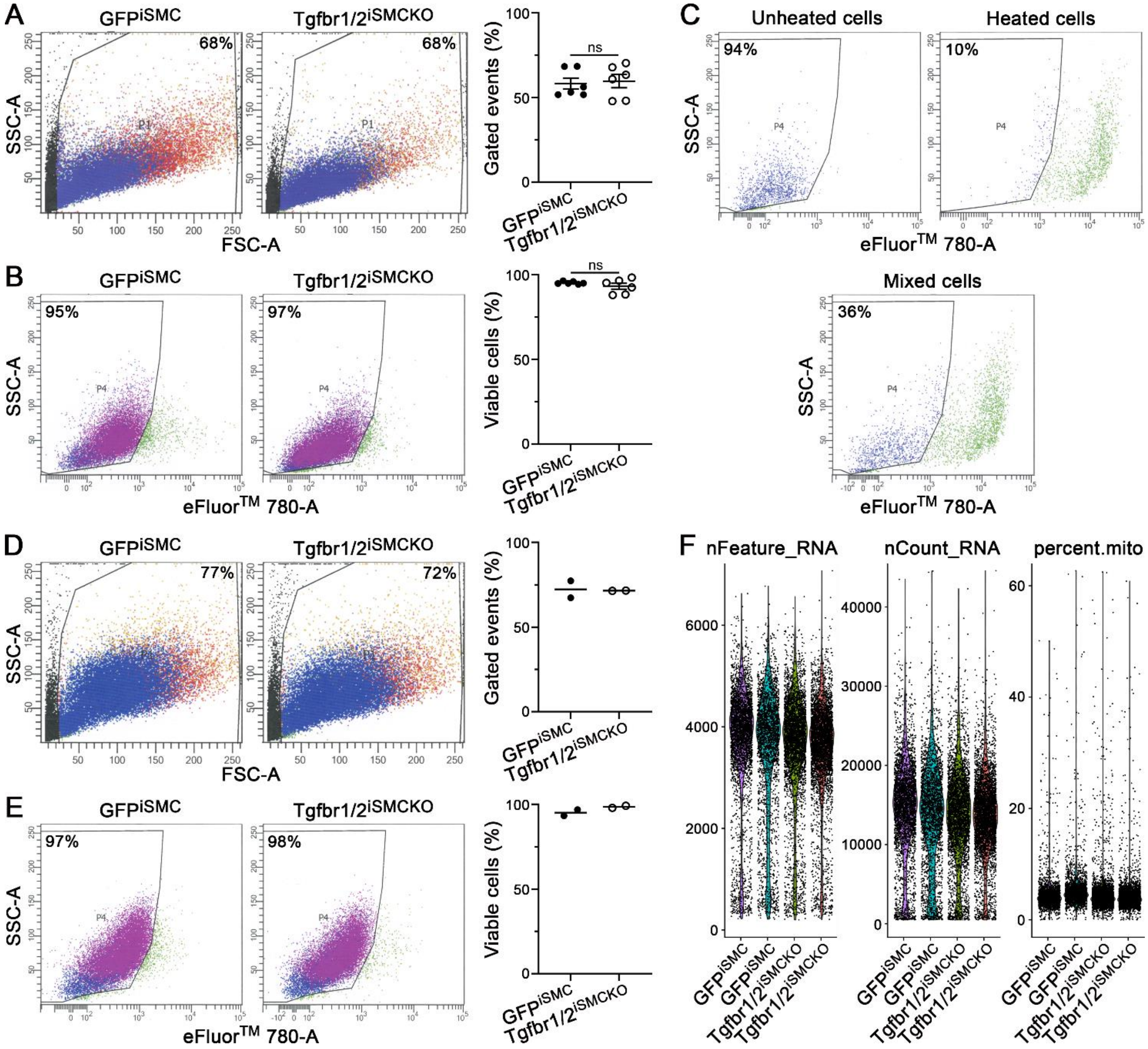


**Supplemental Figure 7: Similar cell viability and RNA quality control metrics in SMCs without and with TGFβ signaling disruption.** Cells were enzymatically isolated from thoracic aorta segments of 12-week-old GFP$^{iSMC}$ and Tgfbr1/2$^{iSMCKO}$ mice for bulk RNAseq analysis (shown in Figure 6A) and were (**A**) distinguished from debris by side (SSC-A) vs. forward (FSC-A) scatter followed by (**B**) selection of viable cells via exclusion of cell impermeant eFluor$^{TM}$ 780 dye. Data are shown as individual values with mean ± SEM bars, *n* = 6, *P* value not significant (ns) by t-test. (**C**) Gating to differentiate viable from injured/dying cells was determined from analysis of distal descending thoracic aorta cells enzymatically isolated at 4 °C and either kept on ice (unheated), heated to 75 °C for 5 minutes (heated), or a mixture of unheated and heated cells (mixed). Alternatively, cells were enzymatically isolated from thoracic aortas of 12-week-old GFP$^{iSMC}$ and Tgfbr1/2$^{iSMCKO}$ mice for single cell RNAseq analysis (shown in Figure 6C) and were (**D**) distinguished from debris by side vs. forward scatter followed by (**E**) selection of viable cells. Data are shown as individual values with mean, *n* = 2, statistical analysis not performed. (**F**) Violin plots for number o f genes per cell (nFeature_RNA), number of total reads per cell (nCount_RNA), and percent mitochondrial reads per cell (percent.mito) prior to filtration and normalization of gene expression results reveal a similar proportion of outliers among experimental groups and replicates.

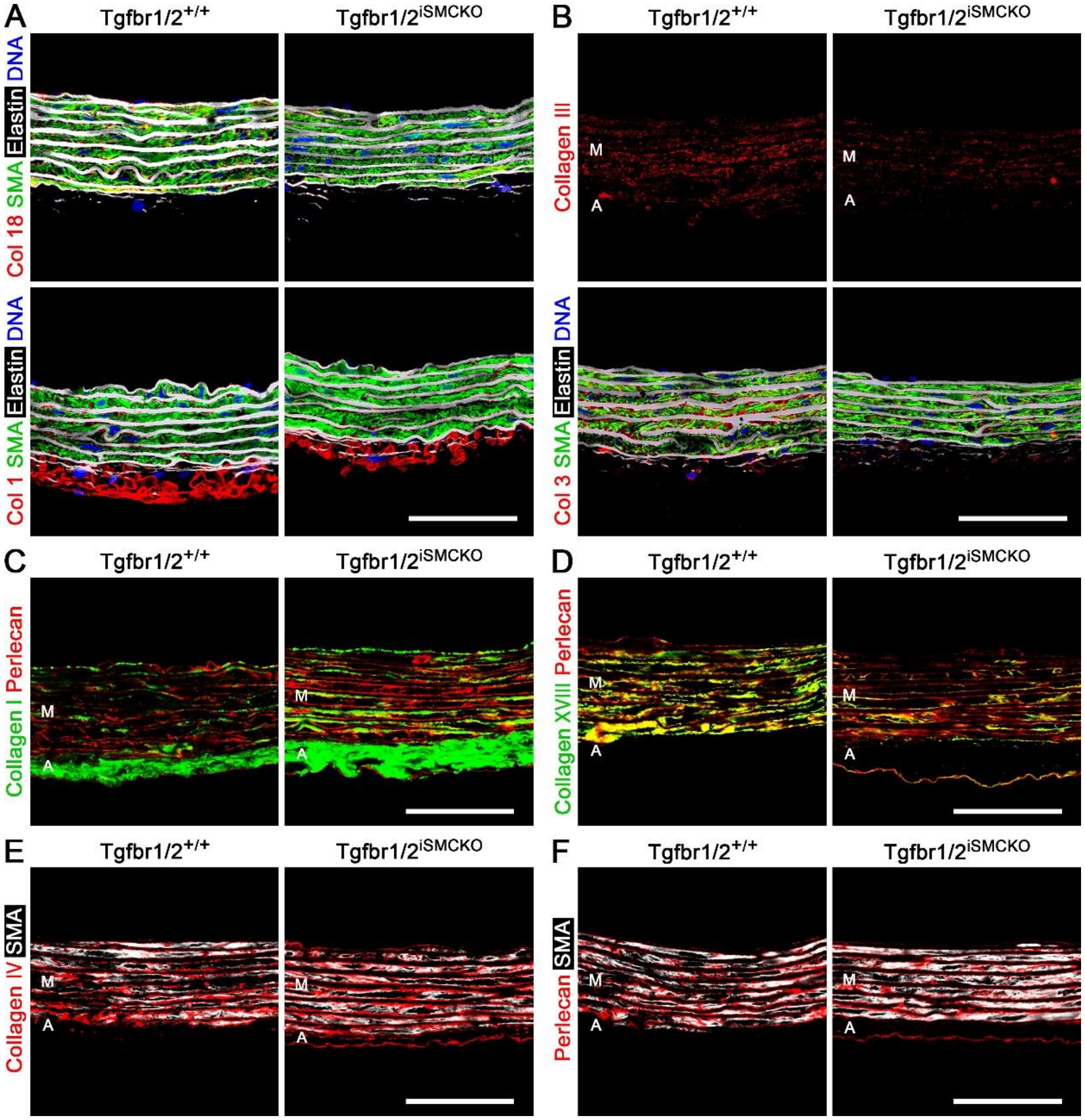


**Supplemental Figure 8: ECM protein expression 1 week after TGFβ signaling disruption in SMCs of mature aortas.** Ascending aortas of 12-week-old Tgfbr1/2$^{+/+}$ and Tgfbr1/2$^{iSMCKO}$ mice at 7 days after starting tamoxifen were analyzed by confocal microscopy. (**A**) Presence of collagen XVIII (Col 18, red) or I (Col 1, red) overlaid with smooth muscle α-actin (SMA) for SMCs (green), AF633 hydrazide for elastin (white), and DAPI for nuclei (blue). The images of collagen XVIII and I alone are shown in Figure 8. (**B**) Presence of collagen III (Col 3, red) alone or overlaid with smooth muscle α-actin for SMCs (green), AF633 hydrazide for elastin (white), and DAPI for nuclei (blue). (**C**) Expression of collagen I (green) overlaid with perlecan (red). (**D**) Presence of collagen XVIII (green) overlaid with perlecan (red) demonstrating co-localization (yellow). (**E**) Presence of collagen IV (red) overlaid with smooth muscle α-actin (white) and (**F**) presence of perlecan (red) overlaid with smooth muscle α-actin (white) showing basement membrane proteins surrounding SMCs. Formalin-fixed, paraffin-embedded sections (A, B) and frozen, OCT-embedded sections (C, D, E, F) of pressure-fixed specimens. The media (M) and adventitia (A) are identified by the presence or absence of elastic laminae (the adventitia is inadvertently trimmed during procurement in some specimens). Scale bars: 50 μm.

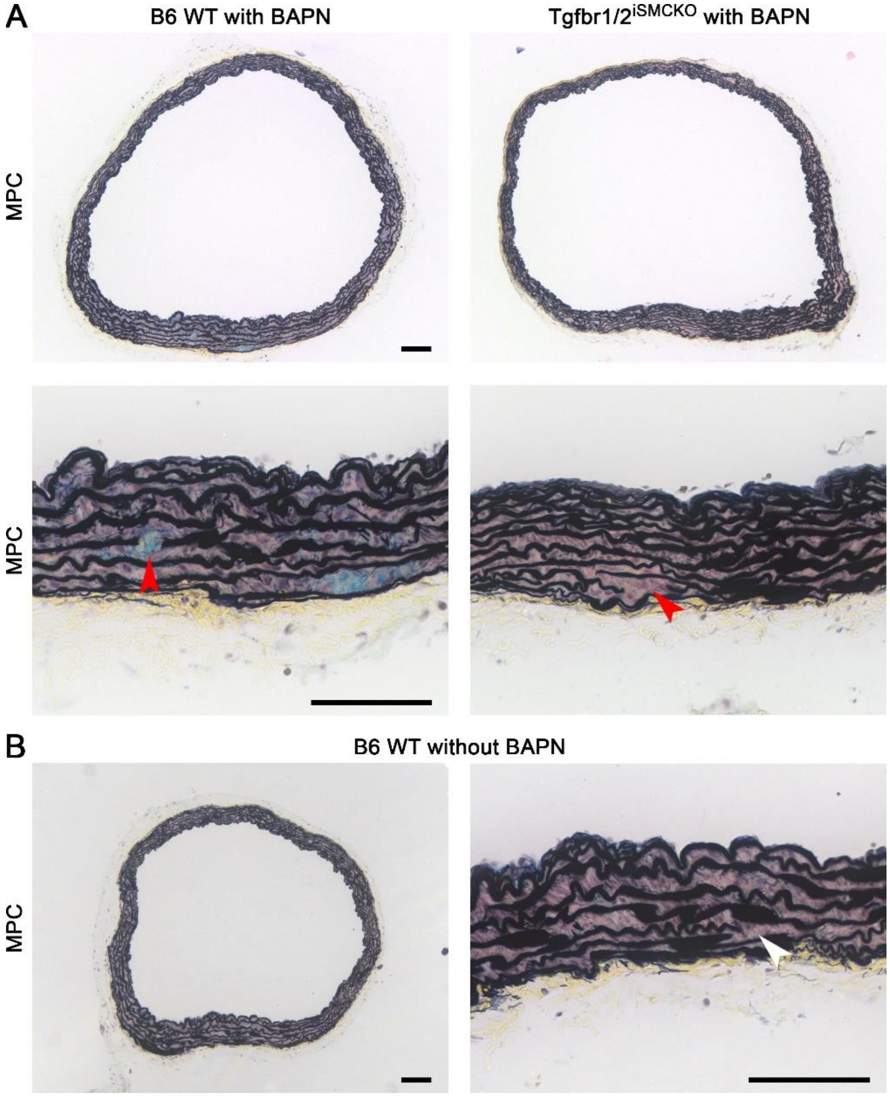


**Supplemental Figure 9: Histological appearance of aortas after short-term BAPN exposure.** Nine-week-old B6 WT and Tgfbr1/2$^{iSMCKO}$ mice were given BAPN at 150 mg/kg/d for 14 days (the mutant animals also received tamoxifen for 5 days starting at 11 weeks of age) and ascending aortas were analyzed at 12 weeks of age when no hemorrhagic lesions were noted macroscopically. (**A**) Transverse sections of formalin-fixed, paraffin-embedded specimens were analyzed with Movat's pentachrome (MPC) stain showing largely unremarkable mural architecture, except for a few focal breaks in elastic fibers associated with mild glycosaminoglycan accumulation or cellular/nuclear hypertrophy (red arrows). (**B**) Ascending aortas from B6 WT control mice without BAPN exposure at 12 weeks of age also displayed infrequent gaps of elastic fibers (white arrow) representing either breaks or lamellar fenestrations since the surrounding ECM and SMCs appeared normal. Scale bars: 100 µm.
